# Regulation of a single Inositol 1-Phosphate Synthase homeolog by HSFA6B contributes to fiber yield maintenance under drought conditions in upland cotton

**DOI:** 10.1101/2022.06.10.495687

**Authors:** Li’ang Yu, Anna C. Nelson Dittrich, Xiaodan Zhang, Venkatesh P. Thirumalaikumar, Giovanni Melandri, Aleksandra Skirycz, Kelly R. Thorp, Lori Hinze, Duke Pauli, Andrew D.L. Nelson

## Abstract

Drought stress substantially impacts crop physiology resulting in alteration of growth and productivity. Understanding the genetic and molecular crosstalk between stress responses and agronomically important traits such as fiber yield is particularly complicated in the allopolyploid species, upland cotton (*Gossypium hirsutum*), due to reduced sequence variability between A and D subgenomes. To better understand how drought stress impacts yield, the transcriptomes of 22 genetically and phenotypically diverse upland cotton accessions grown under well-watered and water-limited conditions in the Arizona low desert were sequenced. Gene co-expression analyses were performed, uncovering a group of stress response genes, in particular transcription factors GhDREB2A-A and GhHSFA6B-D, associated with improved yield under water-limited conditions in an ABA-independent manner. DNA affinity purification sequencing (DAP-seq), as well as public cistrome data from Arabidopsis, were used to identify targets of these two TFs. Among these targets were two lint-yield associated genes previously identified through genome-wide association studies (GWAS) -based approaches, *GhABP-D* and *GhIPS1-A*. Biochemical and phylogenetic approaches were used to determine that *GhIPS1-A* is positively regulated by GhHSFA6B-D, and that this regulatory mechanism is specific to Gossypium spp. containing the A (old-world) genome. Finally, a SNP was identified within the GhHSFA6B-D binding site in *GhIPS1-A* that is positively associated with yield under water limiting conditions. These data lay out a regulatory connection between abiotic stress and fiber yield in cotton that appears conserved in other systems such as Arabidopsis. This regulatory mechanism highlights how sub-genome dynamics contribute to phenotypic stress-response plasticity in cotton.

## Introduction

Upland cotton (*Gossypium hirsutum* L.) is the world’s top renewable textile fiber, supporting a multibillion-dollar industry with a global production of 120.2 million bales of cotton (∼26 million metric tons, USDA, 2022). It is a major economically important crop for the United States and for Arizona, where upland cotton is planted on ∼50,000 ha (USDA 2021), mainly in the semi-arid environment of the low desert using surface irrigation to complement limited precipitation. Cotton productivity in semi-arid areas of the Southwestern United States is severely threatened by global climate change. Increasing climatic variability is responsible for hotter summers, with day and night temperatures far above the thermal optimum (30/22◦C) for the crop, and lower and erratic rainfall patterns which expose the crop to an increasing risk of drought (Alizadeh et al., 2020). Therefore, revealing the physio-genetics mechanisms that regulate cotton’s response to arid conditions is of primary interest. Specifically, this information can be leveraged for the development of new elite cotton cultivars with improved adaptation to hotter and drier climatic conditions that are predicted in the near future.

In addition to being a critical fiber crop, cotton serves as an excellent model polyploid system for studying the impacts that interspecific hybridization has had on agronomic traits. The cotton species predominantly cultivated for fiber production, *G. hirsutum* (upland cotton) and *Gossypium barbadense* (Pima cotton), are New World allotetraploids believed to have formed ∼ 1-2 million years ago from a transoceanic hybridization of an A genome diploid originating from Africa or Asia (e.g., *Gossypium arboreum*, tree cotton) and a D genome diploid from Central or South America (e.g., *Gossypium raimondii*; (Wang et al., 2012; Chen et al., 2007). This unique combination of homeologous gene pairs in the allotetraploids resulted in superior fiber yield and quality over diploid progenitors that has since undergone additional selection in both *G. hirsutum* and *G. barbadense*. Despite high levels of sequence conservation and collinearity between these two species and their diploid progenitors, an altered epigenetic landscape, as well as homeolog expression divergence, have contributed to *G. hirsutum*’s capacity to maintain lint yield under a wide range of environments (Peng et al., 2022; Li et al., 2021; Chen et al., 2020; Pan et al., 2020) Like other crops, drought tolerance in cotton involves complex signaling pathways and transcriptional networks orchestrated by a number of transcription factors (TFs) and signaling proteins (Mahmood et al., 2019; Takahashi et al., 2020; Shinozaki and Yamaguchi-Shinozaki, 2007). These regulatory proteins are typically upstream of, or transcriptionally interconnected with, the genes necessary for coping with the stress and maintaining crucial metabolic pathways. Examples of typical regulatory targets include genes encoding for enzymes related to the production of protective metabolites, transporters, chaperones, and lipid biosynthesis proteins (Malhotra and Sowdhamini, 2014; Singh and Laxmi, 2015; Gupta et al., 2020). In cotton, multiple transcription factor families, such as *GhNACs*, *GhDREBs*, *GhERFs*, and *GhWRKYs*, have been associated with the drought stress response (Huang et al., 2009; Chu et al., 2015; Huang et al., 2013; Ma et al., 2017) Additionally, Genome Wide Association Studies (GWAS)-based approaches have identified numerous candidate genes related to lint yield, fiber quality, and other agronomically important traits (Fang et al., 2014; Sun et al., 2021; Ma et al., 2018). Indeed, while germplasm exists with the ability to maintain fiber growth and quality under heat and drought conditions, little is known about how these traits arose in the cotton genome and the regulatory factors that connect them.

In this study, transcriptome sequencing was performed for 22 upland cotton accessions grown in the Arizona low desert and exposed to both well-watered and water-limited conditions. Phenotypic and metabolomic data were integrated into a co-expression network analysis to identify genes associated with improved yield under water-limited conditions. DAP-sequencing was performed on two transcription factors, *GhDREB2A-A* and *GhHSFA6B-D*, shown to have the highest association with lint yield and stress response genes across the panel. Using transcriptomic, biochemical, and phylogenomic approaches, *GhHSFA6B-D* binds to and positively regulates the lint yield-associated gene, *GhIPS1-A*, and that the associated *GhHSFA6B-D* regulatory element is only present in the A sub-genome and A-genome diploid progenitors. We also identify a lint yield-associated single nucleotide polymorphism directly adjacent to this *GhIPS1-A* regulatory element that influences GhHSFA6B-D binding and appears to suggest additional selection at this locus during domestication.

## Methods and materials

### Plant materials and experimental design

Plant growth conditions have been described in (Melandri et al., 2021), where these same cotton accessions were examined for their metabolite profiles in response to drought stress over a two-year field experiment. In brief, a panel of 22 upland cotton (*Gossypium hirsutum* L.) accessions (**Supplementary Table 1**) were grown at the University of Arizona Maricopa Agricultural Center (MAC) in Maricopa, AZ, United States during the summer of 2019. The accessions were arranged in a randomized incomplete block design with half of the plots experiencing normal irrigation (well-watered, WW) and the other half treated with water-limited (WL) condition, starting when 50% of the plots were at first flower, and consisting of approximately half of the WW irrigation amount. Leaf tissue was harvested ∼50 days later at the boll development stage and used for metabolomics (details in (Melandri et al., 2021)) and transcriptomics analyses. For RNA extraction, two biological replicates, comprised of 10 leaf discs (0.64 cm in diameter) of the upper-most expanded leaf derived from five randomly selected plants (two leaf discs from each plant) in each plot were collected within a single day during a time window of 3 hours (11:00-14:00). Leaf discs were stored in a 2 mL Eppendorf tube containing 1.5 mL of RNAlater (Fisher #AM7021) and stored on ice before being transferred from field to lab where they were then stored in a –80°C freezer before further processing steps.

### Metabolite and phenotypic data measurement

All details on the procedures used to collect/determine the phenotypic trait data and metabolite data used in this study can be found in Melandri et al., (2021). In brief, cotton fiber quantity and quality traits included lint yield (grams/plot), micronaire (Mic, units of air permeability), upper half mean length (UHM, inches), length uniformity (UI, percent), strength (Str, grams per tex) and elongation (Elo, percent). Reflectance-based vegetation indices (VIs) included normalized difference vegetation index (NDVI), carotenoid reflectance index (CRI), scaled photochemical reflectance index (sPRI), and the ratio between the water index (WI) and NDVI (WI/NDVI). The levels of 451 metabolites were determined by untargeted gas chromatography-mass spectrometry (GC-MS, 27 metabolites) and liquid chromatography-mass spectrometry (LC-MS, 424 metabolites). Best Linear Unbiased Estimators (BLUEs) of each phenotypic trait and metabolite value were generated for each cotton accession before being used for downstream statistical analyses.

### RNA-seq library construction

Extraction of RNA from leaf disks was performed using methods from previous studies (Pang et al., 2011; Wu et al., 2002). Hot borate buffer was prepared containing 0.2 M sodium borate pH 9.0, 30 mM EGTA, 1% SDS, and 1% sodium deoxycholate. Just before use, PVP-40, NP-40, and DTT were added to the hot borate buffer to a final concentration of 2%, 1%, and 10 mM, respectively. To extract RNA, a total of approximately 0.75 ml hot borate buffer per 50 mg tissue (∼5 cotton leaf disks) was used. Buffer was heated to 80 °C, and 250 hot buffer was added to 5 leaf disks. Tissue was ground in the hot buffer in a mortar and pestle, then 15 μl of 20 mg/ml Proteinase K (Roche #46950800/Sigma #3115887001) per sample was added and the tissue was ground again. A final 500 ul hot buffer was added before grinding a final time. Lysate was added to a Qiashredder column (Qiagen #79656) and centrifuged at 13,000 x g for 1 minute. Flow through was added to 0.5 volumes of 100% ethanol. The mixture was used as input for RNA cleanup using the RNeasy kit (Qiagen #74104) according to the manufacturer’s instructions. Samples were eluted with RNase-free water. RNA-seq libraries were then generated from mRNA-enriched samples using the Amaryllis Nucleics Full Transcript library prep kit (YourSeq Duet, https://amaryllisnucleics.com/kits/duet-rnaseq-library-kit).

### Transcriptome sequencing and data processing

Each of the 22 *G. hirsutum* accessions treated under both the WW or WL conditions were sequenced using Illumina HiSeqTM 2500 (San Diego, CA) paired-end libraries. Trimmomatic was used to trim adapters and low-quality reads (Bolger et al., 2014). Further, reads were aligned to the G. *hirsutum* reference genome (Ghirsutum_527_v2.0, accession ID: VKGJ01000000, acquired from Phytozome (Chen et al., 2020) , using RMTA v2.6.3 pipeline (Peri et al., 2020) with default parameters and further quantified reads mapped back to each locus using FeatureCounts (parameter: Multimapping; reads: counted; Multi-overlapping reads: counted) and then transformed into length normalized Transcripts Per Kilobase Million (TPM) by custom R scripts (Liao et al., 2014) The raw feature counts for each transcript across 88 samples were normalized by DESeq2 (Love et al., 2014). To test the reproducibility of replicates, we used the Pearson correlation of normalized counts between each set of replicates. Replicates with R^2^ < 0.8 and *P* > 0.05 between samples were removed, leading to only 21 accessions being further examined. Further, pair-wise comparison of TPM values under WL and WW conditions for each accession was performed to identify respective accession-specific DEGs (model: ∼entry + treatment + entry : treatment). For each accession, significantly DEGs under WL condition were classified using thresholds adjusted p-value < 0.01, Log_2_FC > 1 (upregulated) or < 1 (downregulated).

### Variants calling and phylogenetic analysis

Variant calling was conducted with the GATK4 pipeline using the haplotypecaller function (Brouard et al., 2019)The average mapping depth (DP) was calculated across 84 samples as screening cutoff which is equal to 4.12. Then we filtered out variants with sites with DP < 3, depth by quality (DQ) < 2, genotype quality (GQ) < 20, and minor allele frequency < 0.025 by vcftools (Danecek et al., 2011). Phylogenetic relationships were inferred using the IQ-TREE pipeline with filtered SNPs (Nguyen et al., 2015). Briefly, variants calling files (VCFs) of all samples were transformed into phylip format by vcf2phylip tools, a maximum-likelihood tree was constructed (parameter: -nt AUTO -m MFP, Ortiz et al., 2019) and plotted by Figtree (http://tree.bio.ed.ac.uk/software/figtree/). The 21 accessions were subdivided into different groups based on phylogenetic, geographic, and historical breeding information obtained from USDA-GRIN (https://www.ars-grin.gov). A publicly available deep-sequenced (∼20**×**) WGS dataset containing 17 out of 21 accessions used for field experiment was obtained from the NCBI SRA (**Supplemental Table 1**) to identify genome-wide variants. The reads were trimmed by trimonmatic (default settings, (Bolger et al., 2014)) and mapped to the reference genome using bwa-mem along with retention of unique mapped reads by picard (MarkDuplicates -remove = TRUE) for variants calling, The haplotypecaller calling was performed by GATK4 and filtered by VCFtools (--max-missing 0.9, --maf 0.05, minGQ 20, minQ 200, minDP 5, (Danecek et al., 2011)). The filtered variants were annotated using VEP (default settings, (McLaren et al., 2016)) to classify variations to coding regions, promoter, and inter-genic regions.

### Principal component analysis (PCA)

To estimate the strength of treatment effects and the impact of phylogenetic relatedness on transcriptomic and metabolomic profiles, a principal component analysis (PCA) was performed using normalized read counts of 3,768 genes associated with the top 10% of transcriptomic variance or using plotPCA function of DESeq2 (Love et al., 2014), and normalized Z-score of 451 metabolites using “factoextra” R package (https://cran.r-project.org/web/packages/factoextra). Contribution rates for each component were calculated using plotPCA. The first three components from transcriptome and metabolites were visualized in a three-dimensional PCA using Cubemaker (https://tools.altiusinstitute.org/cubemaker/).

### Weighted gene co-expression network analysis

Gene co-expression network was constructed using the WGCNA R package to classify gene expression modules and explore module-trait relationships (Langfelder and Horvath, 2008). Genes with both a high median absolute deviation (MAD) score (top 10%) and genes with high expression (average TPM across all samples > 1) were retained for expression module classification. The pickSoftThreshold function was used to identify the optimal soft power threshold (= 7) at which R^2^ surpassed 0.85 and no further improvements in mean connectivity (module size) were observed, as performed previously (Zhu et al., 2018). Block-wise modules were constructed using the following parameters (power = 7, maxBlockSize = 5000, TOMType = “unsigned”, minModuleSize = 30, reassignThreshold = 0, mergeCutHeight = 0.2). Trait-module relationships were derived using the “modTraitCor” function in WGCNA. Modules that displayed a high trait correlation (R^2^ > 0.6, *p* < 0.05) were selected for further analysis. To investigate modules with high module-trait membership, only co-expressed genes with strong connectivity (weight score > 0.1) were retained for downstream analysis.

### Functional characterization of trait-correlated module

Gene functional annotations of filtered genes (weight score > 0.1) in lint yield-correlated module were obtained from the cotton functional genome database (CottonFGD) to perform enrichment of gene ontology (GO) and KEGG pathways (https://cottonfgd.org/(Zhu et al., 2017)) using Fisher Exact test (*P* < 0.01). Transcription factors (TFs), transcription regulators (TRs), and protein kinases within this module were classified using iTAK (Zheng et al., 2016). Reported protein-protein interactions (PPIs) among genes within the same module were screened using the STRING database (Szklarczyk et al., 2017) (confidence level = 0.6, interaction source: database, experiments). To identify the upstream master TF(s) that may bind to genes within the lint yield module, the enrichment of consensus TF motifs was tested using the 2-Kbp upstream region of the 866 co-expressed genes within the module. The enrichment test was performed by the Analysis of Motif Enrichment (AME) pipeline (McLeay and Bailey, 2010) with the *Arabidopsis* DAP-seq profiles of (O’Malley et al., 2016) as the consensus motif database (parameters: --scoring avg --method fisher --hit-lo-fraction 0.25 --e-value --kmer report-threshold 10.0, cutoff: TP values > 3, *P*-value < 0.001).

### DNA affinity purification sequencing (DAP-Seq) library construction

The four TFs were selected as hub genes to construct DAP-seq libraries, including two *GhDREB2A* (*Gohir.A13G021700* and Gohir.D13G022300) and two *GhHSFA6B* (*Gohir.D08G072600* and *Gohir.A08G064100*). DAP-Seq assay was carried out as described by published protocol (Bartlett et al., 2017). The NEB Next® DNA Library Prep Master Mix set for Illumina kit (NEB #E6040S) was implemented to prepare the DAP-Seq gDNA library. The pIX-HALO Vector (cat#G184A, Promega) was used to fuse the *GhDREB2A* and *GhHSFA6B* into the HaloTag. We further used the TNT SP6 High-Yield Wheat Germ Protein Expression System (L3260, Promega) to express the four TFs-HaloTag fusion protein. Magens HaloTag Beads (G7281, Promega) was used to purify the fusion protein. The fusion protein and 500Lng of library DNA were co-incubated in 40 μl PBS buffer for 1.5 hours shaking in a cold room. The beads were washed with 200 μl PBSL+LNP40 (0.005%) for 5 times. The supernatant was discarded and an aliquot of 25 μl of elution buffer was added. Finally, beads were incubated at 98L°C for 10 min to elute DNA fragments. According to the fragment size of the library, the DAP-Seq library concentration for a given read count was measured.

### DAP-Seq sequencing and peak analysis

DAP-seq libraries were sequenced by Illumina short reads platform (Single end: 150 bp) with a total of expected 40 million reads for each protein to be sequenced. These reads were trimmed by TrimGalore (Babraham et al., 2015) and then mapped to the reference genome (*TM-1*: Ghirsutum_527_v2.0, Phytozome accession ID: VKGJ01000000)(Chen et al., 2020) using bowtie2 and sorted by sambamba. Before peak calling, only the unique mapped reads were retained by sambamba (parameter: -F “[XS] == null and not unmapped and not duplicate”) to prevent false positives due to multiple mapped hits (Tarasov et al., 2015). Furthermore, the MAC2s tool was applied to call peaks (parameter: --keep-dupall -g 2.3-e9) and high confidence peaks were captured using IDR (Padj < 0.05)(Zhang et al., 2008). These high-quality peaks were annotated based on reference gene annotation using the ChIPseeker R package (Yu et al., 2015) . Regulatory regions of a target gene were defined as the 5 Kb upstream sequences before the transcription start sites (TSSs) and the downstream sequences which bearing the longest 5’UTRs among respective isoforms. In particular, only DAP-seq peaks which fell into the regulatory regions were kept as TF targeted genes. The sequences associated with DAP-seq peaks of TFs were harvested as query sequences to perform motif discovery by MEME suite (Bartlett et al., 2017), to identify conserved consensus motif sequences among peak sequences (parameter: -mod zoops -nmotifs 3 -minw 6 -maxw 50 -objfun classic -revcomp -markov_order 0).

### Genomic analysis of TF binding regions

Paralogous genes of the selected lint module genes (*GhABP* and *GhIPS*) were identified using Blastn (Ye et al., 2006) , which aligns the CDS sequences against the reference gene annotation (cutoff: E-value < 10-3, identities > 90%). Those hits been identified were further checked by functional annotation. Further, the promoter and coding regions of these paralogous gene pairs were extracted and then aligned using multiple sequences alignment by Geneious (tool: MUSCLE, iteration: 500) to identify variants (SNPs and INDELs) between paralogous gene pairs, specifically the regions bearing the DAP-seq peak (Kearse et al., 2012). The potential binding sites of two TFs over the peak regions were defined by motif scan (FIMO) using *Arabidopsis* DAP-seq derived consensus motif sequences (Motif ID: ERF48_col_a, HSFA6B_cal_a, HSFA6B_colamp_a) (Bailey et al., 2015).

### *In-vitro* protein expression and electrophoretic mobility shift assay (EMSA)

Protein expression was carried out using the *in vitro* transcription and translation system (TnT™ T7 Quick for PCR DNA, Promega) and the EMSA reaction was carried out following manufacturer’s instruction (odyssey). The vector containing the gene of interest (with flag tag) was PCR amplified to obtain the DNA template. The reaction mixture contained the DNA template, reaction buffer, amino acid mix, and T7 RNA polymerase, following the manufacturer’s instructions. The reaction was incubated at the recommended temperature for a specified duration, and the resulting protein product was subsequently analyzed by SDS-PAGE to confirm expression.

Immunoblot analysis were performed by using standard wet transfer method. The proteins were blotted using nitrocellulose membrane (Sigma Aldrich). Rabbit monoclonal halo tag antibody was used in the (1:5000) dilution (v/v). Horse-radish peroxidase conjugated goat anti-rabbit antibodies were used as secondary antibodies. Blots were developed using Super-signal Chemiluminescent substrate following the manufacturer’s instruction. The blots were imaged using the BioRAD gel imager following the linear curve of detection in the signal.

The interaction between the expressed protein and its putative DNA-binding partner was analyzed using EMSA. A double-stranded DNA probe containing the target binding site was prepared by annealing complementary oligonucleotides labeled with a fluorophore. The binding reaction mixture consisted of the expressed protein, labeled DNA probe, binding buffer, and appropriate cofactors. The reaction mixture was incubated at room temperature for 30 minutes to allow for protein-DNA complex formation. Subsequently, the samples were resolved on a non-denaturing polyacrylamide gel electrophoresis (PAGE) gel with a 6% acrylamide concentration. The gel was run at 120 volts for 3 hours in 1**×** TBE. Following electrophoresis, the gel was visualized using LiCor Odyssey Gel Scanner to capture the mobility shifts indicating protein-DNA complex formation. For competition assays, unlabeled competitor DNA or unlabeled mutated competitor DNA was added to the binding reaction at increasing molar excesses.

## Results and Discussion

### Genetic profiles of upland cotton panels in field experiment

To better understand the molecular mechanisms underlying cotton’s performance under drought conditions, 22 diverse upland cotton accessions from the Gossypium Diversity Reference Set (Hinze et al., 2015, 2016) were evaluated under water-limited (WL) and well-watered (WW) conditions in the field. Leaf tissue from plants at the flowering and boll development stage was collected (two replicates each, ∼ 100 DAS; **Figure 1A**). On average, ∼27 million PE x 150 bp reads were generated for each replicate. Reads were then aligned using the RMTA pipeline (Peri et al., 2020) to the G. *hirsutum* v2.0 reference genome (Chen et al., 2020); (**Supplementary Table 1**). Gene level expression data were counted using *G. hirsutum* gene-level annotations as meta-features in FeatureCounts (Liao et al., 2014; **Supplementary Table,** parameter: -t gene, -g ID). The Pearson correlation coefficient (PCC) of normalized read counts revealed high correlation among replicates (average PCC = 0.904), except for accession “Tipo Chaco” (PCC = 0.61, cutoff: 0.8) (**Supplementary Table 2**). Thus, this accession was discarded from further analyses resulting in 21 accessions for further analyses.

**Figure. 1.**
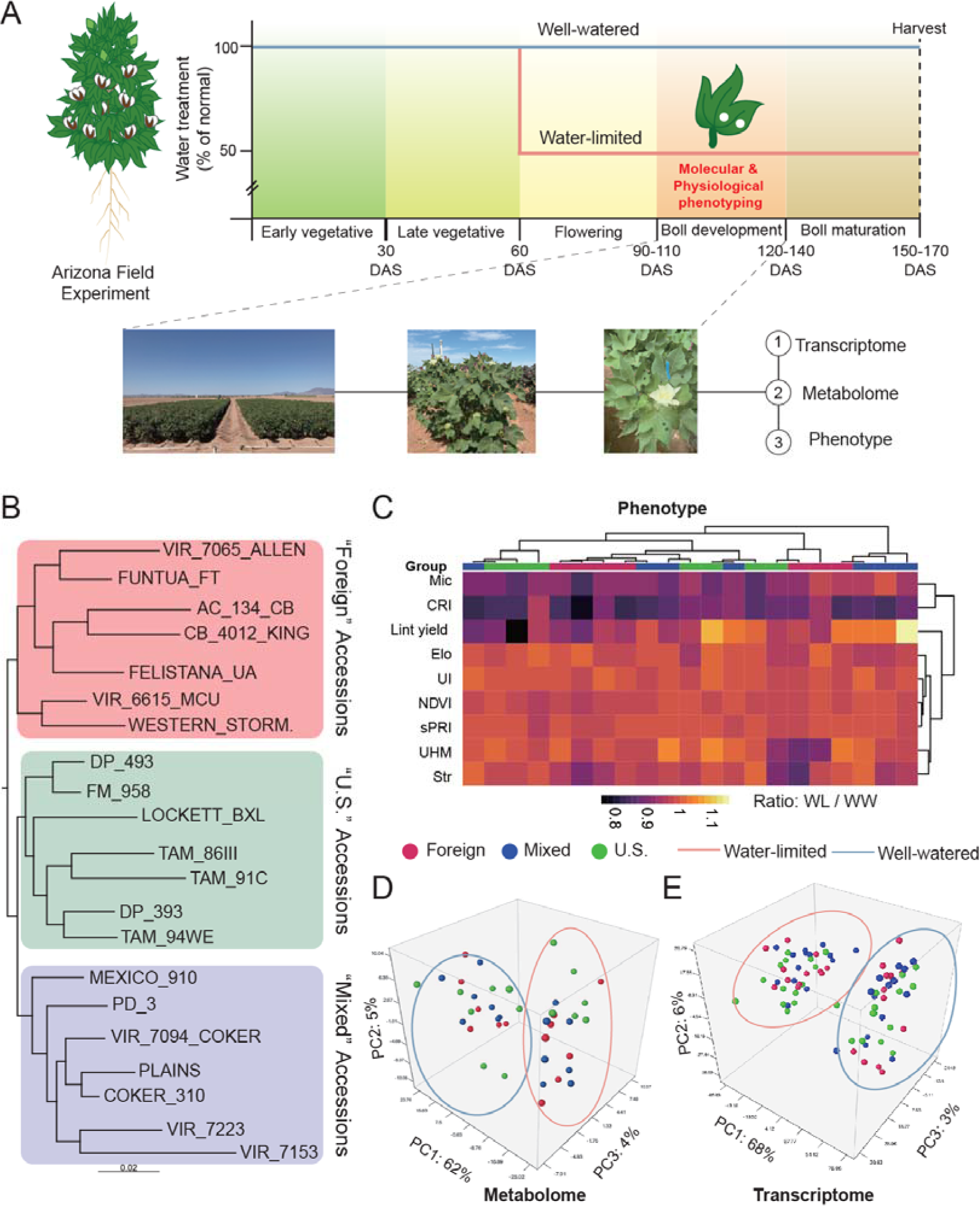
Experiment information and genomic, metabolomic, and transcriptomic profiling of the cotton panel used in this work. **A)** Overview of the watering regime and timing of data collection for the experiment in the 2019 summer field season at Maricopa, AZ. Water levels were reduced to 50% of normal for the treatment plots at the early flowering stage. Metabolomic and transcriptomic data were collected at late flowering/early boll development, with phenotypic measurements taken throughout (See Methods). **B)** A phylogeny of the accessions used in this study based on filtered SNPs derived from the transcriptomic data. The three groups, “Foreign”, “U.S.”, and “Mixed”, reflect historical breeding information obtained from USDA GRIN-Global. **C)** Overview of the phenotype profiles of 21 upland cotton accessions under water-limited (WL) and well-watered (WW) conditions: A cluster heatmap and indicated the ratio of phenotypic data under WL condition relative to the WW condition (WL value / WW value, range: 0.8 -1.2). Group coloring denotes accessions belonging to the three phylogenetic groups from 1B. **D)** Three-dimensional PCA displaying the impact that treatment has on large-scale changes in normalized metabolite content between accessions. The different treatments have been denoted with light red (water-limited) and light blue (well-watered) ovals. **E)** Three-dimensional PCA was used to display the top 10% (*N* = 3768) most variably expressed genes based on normalized read counts, with treatment groups denoted similarly to **1D**.

To determine the relatedness of each of these accessions, these RNA-seq data were used to perform variant calling relative to the reference genome (108,396 bi-allelic SNPs after filtering). These data were used to reconstruct a phylogeny of the 21 accessions, rooted by VIR_7153_D_10, which resulted in three major subgroups (**Figure 1B**). Based on information from USDA GRIN-Global, the first of these three subgroups is referred to as “Foreign”, as this group contains accessions developed primarily outside of the U.S. (**Figure 1B** and **Supplementary Table 1**). The second subgroup consists of “U.S.” accessions of breeding lines and commercial cultivars developed in the U.S. (**Figure 1B** and **Supplementary Table 1**). The last group, “Mixed”, consists of commercial cultivars from the U.S. and improved breeding lines generally developed outside the U.S. (**Figure 1B** and **Supplementary Table 1**). Thus, these 21 accessions serve as a genetically diverse framework to investigate the regulatory mechanisms controlling the physiological responses to drought in cotton.

### Variation in physiological and molecular responses to drought in major subgroups

The phylogenetic groupings were used to determine the degree to which genotype and environment influenced physiological responses to drought in the assembled panel. The effects of genotype (G: Foreign, U.S., and Mixed), environment (E: WL and WW), and G*E interaction on six fiber quality and agronomic traits and four vegetation indices, were examined using two-way ANOVA (**Supplementary Table 3**). Among these traits, three of the four vegetation indices were significantly affected by drought (**Supplementary Table 3;** two-way ANOVA cut-off: *p* < 0.05, **Figure. S1**). The treatment effect was particularly significant in the “Foreign” subgroup (student *t*-test cutoff: *p* < 0.05). By contrast, long-term drought stress brought limited impacts to most fiber quality traits apart from the significantly reduced micronaire (two-way ANOVA: *p* = 0.002) observed in “Mixed” and “Foreign” subgroups (t-test: *p* = 0.015 and 0.047, respectively, **Figure. S1, Supplementary Table 3**). In contrast to treatment effects, pronounced genotypic effects were observed between major subgroups (“Foreign, U.S. and Mixed) for five of the six fiber quality traits but not for the four vegetation indices (**Supplementary Table 3**).

However, intragroup comparisons of responses to drought revealed stark differences. For instance, each subgroup contained one or more accessions that significantly outperformed close relatives when using a ratio of phenotypes from WL relative to WW condition as an indicator, particularly in fiber quality traits (**Figures 1C**). The lack of phylogenetic congruence in observed traits suggests that lint yield and drought associated traits may have been selected differently in even closely related accessions. Importantly, the presence of outperforming accessions implies that this panel may be useful in identifying the factors associated with lint yield and drought stress.

We analyzed metabolite profiling as molecular phenotypes for stress response understand the effects of drought stress. Changes in the metabolome of the panel were measured by re-examining 451 metabolites previously analyzed in Melandri et al., 2021 (27 GC-MS and 424 LC-MS/MS).). A global profiling of the 451 metabolites revealed that the strongest effects were due to drought treatment (PC1, 62%; **Figure 1D**). The second and third principal components explained a further 5.0% and 3.4%, respectively, but the phylogenetic relationships (i.e., genotype) were not clearly separated along either of these axes. A large treatment effect was also observed in the metabolite profiles using a two-way ANOVA **(Supplementary Table 4**). Drought treatment significantly impacted more than 95% of metabolites, but only 5% of the metabolites exhibited genotypic effects; around 25% of metabolites exhibited changes that could be associated with G*E effects (*p* < 0.05, **Supplementary Table 4**). Among the 432 metabolites associated with significant treatment effects, 273 were up-regulated and 159 down-regulated by drought. Among these metabolites are the known glycine and proline (Fang et al., 2015); **Supplementary Table 4**). Thus, these data demonstrate that alteration of specific metabolites associated with drought tolerance (osmoprotectants) is a common response to water deficit stress conditions across our panel.

To test the basis for metabolic changes, we examined transcriptome profiles among these accessions in response to drought stress. Consistent with the observed metabolomic changes, 68% of the variation within the panel could be explained by the irrigation treatment (**Figure 1E**). In addition, neither PC 2 nor PC 3 could be attributed to genotypic differences. Pairwise comparisons of gene expression under the two conditions generated a range of differentially expressed genes (DEGs) for each accession **(Supplementary Table 5)**. We further compared transcriptomes to determine if DEGs were shared between all intragroup accessions, or between all accessions within multiple subgroups in response to drought. Out of the thousands of DEGs in each accession, there were few genes that were shared among all accessions within a subgroup, or between subgroups (**Figure S2A, Supplementary Table 5**). The foreign accessions shared the least group-wide DEGs (up-regulated: 57 and downregulated: 86), whereas the mixed accessions featured the most (up-regulated: 194 and down-regulated: 177) (**Figure S2B**). In contrast, there were 300 DEGs (194 upregulated, 106 downregulated) shared among all examined accessions. Gene ontology (GO) enrichment of these shared up-regulated genes revealed an over-representation of the stress response and phosphorylation signal transduction genes while the shared down-regulated genes were involved in fatty acid biosynthetic processes and transport processes (**Figure S2C**). Thus, these shared DEGs may represent a common set of genes involved in drought stress response.

### Using WGCNA to examine transcriptome-trait connections in response to drought

To determine a relationship between transcriptomic and phenotypic variability, we performed a weighted gene co-expression analysis (WGCNA) to uncover the association of gene networks with phenotypes. For the WGCNA, we incorporated the top 10% most variable genes across the 21 accessions under WL conditions (determined by median absolute deviation [MAD], *N = 4,552* genes). We obtained 22 modules that satisfied a scale-free topology (R^2^ = 0.86; **Figure S3A**) using a soft threshold (Beta = 7) for network construction (**Figure S3B**). These 22 modules displayed clear separation, with only 45 genes (< 0.2 %) unclassified (**Figure S3C**). Using PCC, these 22 well-clustered modules were then correlated to the trait dataset, which consisted of 10 phenotypes and 451 metabolites. Among these traits being tested, five traits (lint percentage, referred to here as lint yield, UI, sPRI, CRI, WI/NDVI) and nine metabolites exhibited significant correlation to 18 modules (**Figure 2, *P* < 0.05**). In particular, lint yield displayed the highest positive correlation to the turquoise module (*r* = 0.68, *p* = 7e-04), whereas terpenoid and carbohydrate metabolites displayed a positive correlation with the salmon module (*P* < 0.05; **Figure 2**).

**Figure. 2.**
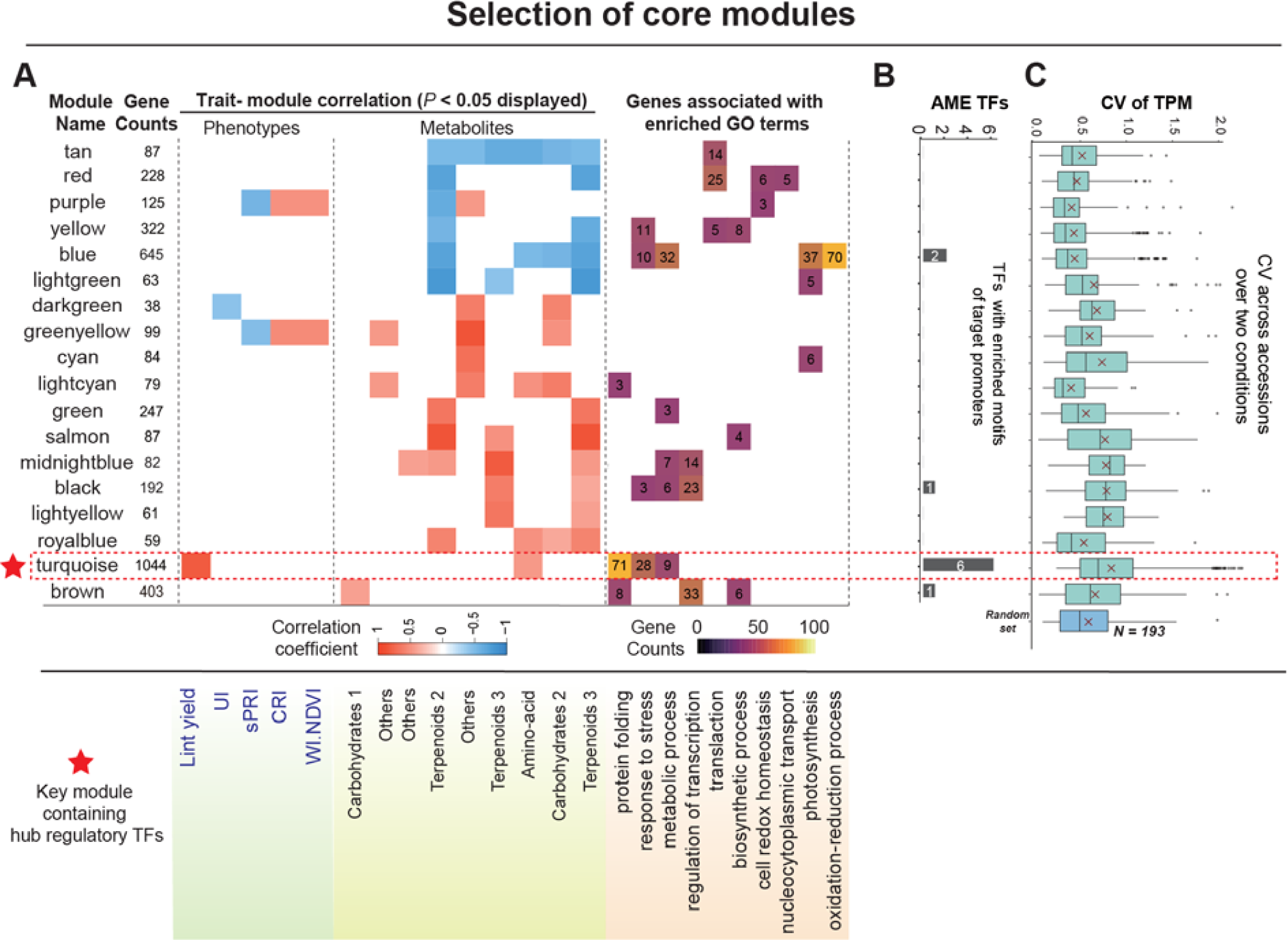
Trait and co-expression association in response to water deficit conditions. **A)** Modules of genes derived from a weighted gene co-expression analysis were correlated with phenotypic and metabolomic traits, with traits passing the significance threshold (*P* < 0.05) shown along the bottom. Modules are named according to color, with the number of genes within each module shown (gene counts). Positive (red hues) and negative (blue hues) correlation values are shown for all significant trait-module correlations with scale bar below. GO term enrichment was performed on all genes within the module, with significant terms shown along the bottom. The number of genes associated with each GO term is shown within each box along with the “purple” to “yellow” scale displayed. **B)** The number of TFs with enriched binding motifs within their respective module (TF and motifs are present in the same module) are shown. **C)** Coefficient of variance (CV) of expression (TPM) was calculated for all genes within each module across all accessions and both conditions. CV was also calculated for a background set of genes based on an average module size (*N* = 193). The red dashed box outlines the “turquoise” module, which was found to be correlated with both lint yield and stress-response genes and was selected as a key module (highlighted with a red star) for further examination.

### Identifying trait-associated functional modules using multiple data integration

Following module-trait correlation, we selected key module(s) for further functional analysis by incorporating the profile of GO enrichment, transcription factor (TF) binding motif enrichment, and gene expression variance (**Figure 2A**). Using Fisher test (*P* < 0.01) for GO enrichment, 14 of the 18 modules exhibited some degree of enrichment for genes involved in biological processes of interest. Notably, the lint yield-associated module contained a high degree of enriched stress response genes (Q-value: 1.6e^-12^) as well as protein folding genes (Q-value: 4.6e^-8^). Also, a large number of photosynthesis-related and redox-related genes (Q-value: 2.3e^-10^ and Q-value: 8.4e^-4^, respectively) were identified within the blue module, negatively correlated with the abundance of multiple metabolites (**Figure 2A**). These data suggest that a co-expression network-based approach may uncover key genes integrating plant development and yield in the response to water limitation.

To further investigate the upstream regulators of those genes associated with enriched GO terms, we took the 2-kb upstream promoter region sequences of all genes (WGCNA weight score > 0.1) in each of the 18 modules to perform motif enrichment analysis using AME (Analysis of Motif Enrichment; (McLeay and Bailey, 2010)). To narrow down the list of possible enriched TFs/motifs, we focused on TF/motif pairs where the TF was also present in the same module and thus displayed similar expression patterns across treatments and genotypes. This approach uncovered four modules with enriched motifs corresponding to 10 different transcription factors, with the lint-yield module (turquoise) containing the most enriched TFs (*n* = 6, AME TFs, **Figure 2B**). Under the hypothesis that significant trait-associated modules might display a more pronounced response to treatment, we assessed expression variation for genes within each module between accessions and conditions. While turquoise module showed a significant increase (*P* = 0.0033) in coefficient of variance (CV) relative to background, and this module contained numerous stress response genes with high CV (**Figure 2C**). In sum, these data indicate that the genes within the “turquoise” module (highlighted with a star in **Figure 2A**), and their associated TFs, may be critical for maintaining yield under water limiting conditions.

### DAP-seq of HSFA6B and DREB2A revealed the module-specific upregulation of their targets in response to drought

Among six TFs in the turquoise module that are associated with yield, four were AME TFs that are predicted to be associated with heat or drought stress based on homology with functionally characterized TFs in *Arabidopsis* and cotton(Huang et al., 2016; Bian et al., 2020; Jacob et al., 2017; Kolmos et al., 2014; Chen et al., 2017; Nakashima et al., 2014; Weltmeier et al., 2009, Figure 3A). We further compared the gene from the enriched GO/KEGG terms (***N = 119***) in this module, as well as seven genes identified in previous GWAS, with lint yield for the presence of TF motifs ((Su et al., 2016a; Sun et al., 2021; Fang et al., 2017)**, Supplementary Table 5**). These four TFs: HSF7, HSF6, HSFA6B, and DREB2A (**Figure 3A**, upper panel) are predicted to bind to a number of stress and heat response genes (**Figure 3A**, bottom panel). Notably, only HSFA6B and DREB2A are predicted to bind to the seven lint-yield associated genes in the module (**Figure 3A**, boxed, and **Supplementary Table 7**). This suggests that the cotton DREB2A and HSFA6B homologs are likely candidates for the observed association between water stress and fiber yield.

**Figure. 3.**
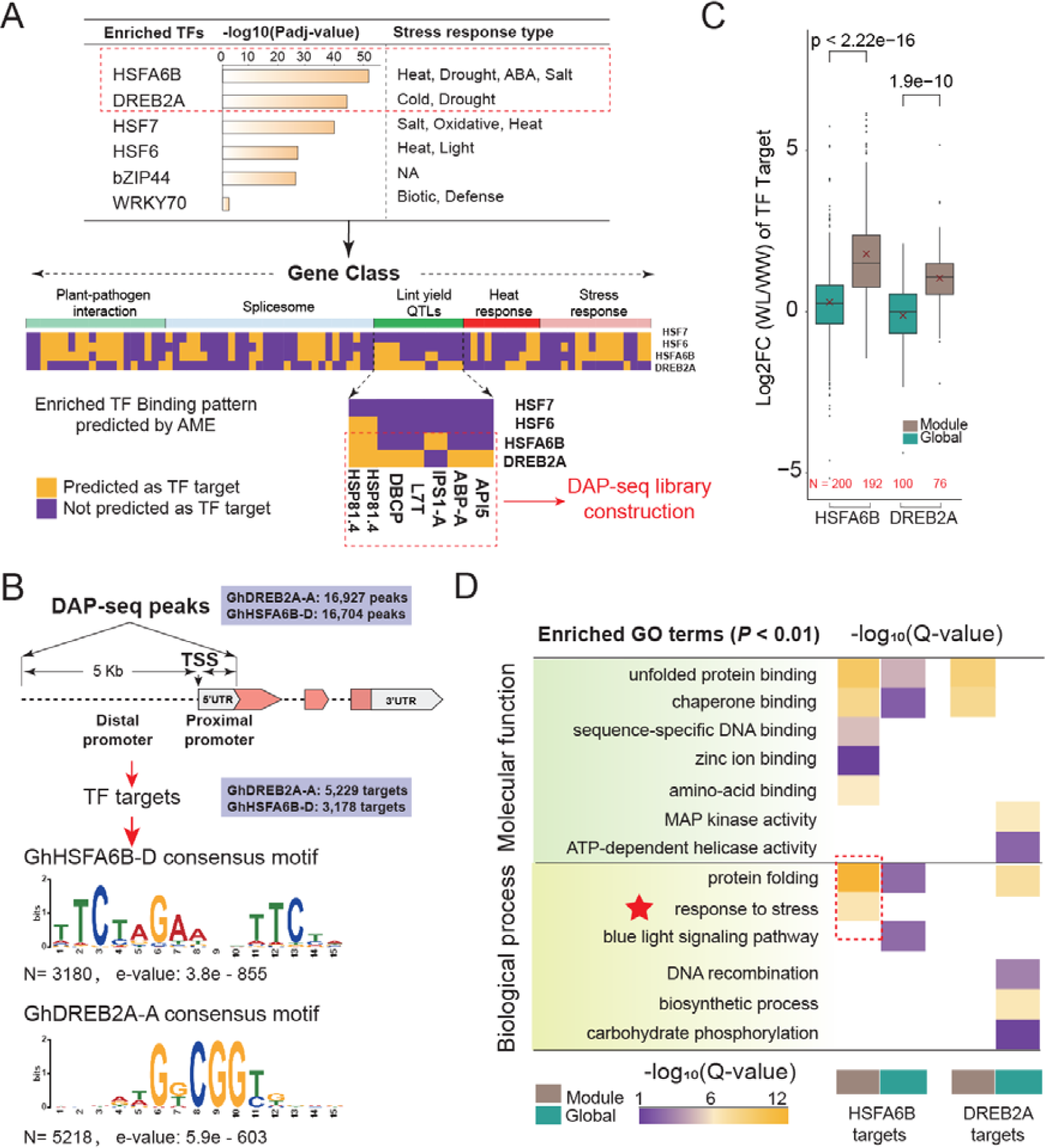
Identifying HSFA6B and DREB2A targets with DAP-seq. **A)** Identity of the seven key hub TFs within the lint yield associated module (Top). For each TF, the -log10 (Adj-P value) level of motif enrichment among co-expressed genes within the module is shown, along with the reported stress response. For the four abiotic stress-associated TFs, their AME predicted binding to the genes with enriched GO terms within the module, as well as five GWAS-identified lint yield genes, is shown (bottom). As GhHSFA6B-D and GhDREB2A-A were both stress response regulators and predicted to bind to at least one of the five lint yield associated genes, they were chosen for DAP-seq library creation (red dashed box). **B)** Illustration of annotating peaks derived from GhHSFA6B-D and GhDREB2A-A DAP-seq data. The top panel highlights DAP-seq peaks within the 5-Kb upstream regions (distal promoters) and 5’UTR regions (proximal promoters) of annotated genes. The bottom panel depicts the two consensus motifs identified from 3,180 peaks (GhHSFA6B-D) and 5,218 peaks (GhDREB2A-A) that passed stringency filters. **C)** Comparison of the log_2_ fold change of transcript abundance (WL relative to WW) for GhDREB2A-A and GhHSFA6B-D DAP-seq targets between genes in the lint yield associated module (192 GhHSFA6B-D targets and 76 GhDREB2A-A targets) and genes selected from transcriptome-wide through random sampling (*N* values shown below). Pairwise significance was performed using a paired Student’s *t*-Test. **D)** Comparison of enriched GO terms among genes bound by GhHSFA6B-D and/or GhDREB2A-A within the lint yield module and targets from a transcriptome-wide group of expressed transcripts. The level of enrichment for each GO term is reflected by -log_10_- Qvalue, with higher levels of enrichment corresponding to larger values. The “star” logo highlights the “stress response” enriched GO term.

Given their association with fiber yield and drought stress, these two TFs likely regulate downstream stress responsive genes. To test this, we performed DNA affinity purification sequencing (DAP-seq) using cotton DREB2A and HSFA6B and examined the DNA-binding ability of the homoeologs from both subgenomes (e.g., GhDREB2A-A and GhDREB2A-D). More than 40 million (DREB2A) and 37 million (HSFA6B) single-end reads (SE: 150 bp) per replicate enabled the identification of more than 20,000 peaks with high reproducibility for each sample after peak calling. For each TF, despite both being expressed in an *in vitro* wheat germ system (**Figure S4**), only a single homoeolog, namely, GhHSFA6B-D, derived from the D subgenome (Gohir.D08G072600) and GhDREB2A-A, derived from the A subgenome (Gohir.A13G021700), could bind to DNA above background. Using Irreproducibility Discovery Rate as a cutoff (IDR, *P* < 0.05), a total of 16,927 and 16,704 high confidence peaks were identified between replicates of GhDREB2A-A and GhHSFA6B-D **(Figure 3B, Supplementary Table 8**). After filtering for GhHSFA6B-D and GhDREB2A-A peaks in the 5-kb upstream region of annotated coding genes (distal promoter), as well as peaks within the 5’ untranslated regions (UTRs) of coding genes (proximal promoter), a total of 5,229 and 3,178 genes were identified as likely regulatory targets of GhDREB2A-A and GhHSFA6B-D, respectively (**Figure 3B, Supplementary Table 9 and 10**). Genomic sequences associated with these peaks were further processed by motif analysis (MEME suite; Bailey et al., 2015) to identify binding sites and test the levels of conservation of consensus motifs. As expected, both target sequences of the two TFs revealed high-level conservation (E-value: 3.8^e-855^ across 3180 peaks and 5.9^e-603^ among 5218 peaks; **Figure 3B**), compared to core HSFA6B and DREB2A binding elements (HSEs and DREs) from other species (Scharf et al., 2012; Sakuma et al., 2006; Nishizawa et al., 2006). Interestingly, although our AME TF approach only predicted a GhHSFA6B-D binding motif in the lint yield-associated gene *GhIPS1-A*, (Myo-inositol-phosphate synthase protein), we observed DAP-seq peaks for GhHSFA6B-D binding to both *GhIPS-A* and *GhABP-A* (**Figure S5**). In addition, we observed GhHSFA6B-D peaks in the promoter region of both *GhDREB2A-A* and *GhDREB2A-D* homeologs (**Figure S5)** but did not observe reciprocal GhDREB2A-A peaks in the promoter of *GhHSFA6B-D*, suggesting that GhHSFA6B-D may act as a regulator of *GhDREB2A-A*. Together, these data suggest that GhHSFA6B-D and GhDREB2A-A may regulate expression of stress responsive pathway genes under drought in cotton.

To determine if GhHSFA6B-D and GhDREB2A-A might have a specific impact under stress on genes found within the lint yield module, we profiled transcriptome changes between treatments. A comparison of average log_2_FC (WL relative to WW) for in-module targets of GhHSFA6B-D or GhDREB2A-A revealed a significant up-regulation relative to global putative TF targets (*P* < 0.001, **Figure 3C**). The analysis of GO enrichment (*P* < 0.01) for these downstream target genes found that only the genes bound by GhHSFA6B-D in the module were enriched for stress response terms (-log_10_(Q-value) > 8), whereas both GhHSFA6B-D and GhDREB2A-A exhibited in-module specificity to chaperone/protein folding related genes (**Figure 3D**). These data suggest that while both TFs display a high degree of regulatory connections between fiber yield and stress responses, GhHSFA6B-D may be a major regulator.

To better visualize the interactions between these two TFs and other genes within the lint yield module, including each other, we synthesized our data into a regulatory network using Cytoscape (**Figure 4A**). To simplify this network, 866 genes in this module were reduced to a subset of 126 core genes by selecting genes with enriched GO terms, KEGG pathways, annotated as TFs or lint yield associated genes, and those highly correlated with lint yield (trait-correlated genes; **Figure 4A**, see key). Trait-correlated genes are those with high module membership (MM), which is derived from the correlation between gene expression and the eigenvalue (first principal component) of the lint yield module, and gene significance (GS), a term reflecting the correlation between gene expression and lint yield (**Figure 4B**, orange circles). These target genes are those whose expression change across the 21 accessions is most likely to explain variation in lint yield in this panel. To further develop this network, we integrated the co-expression information derived from WGCNA, TF-target binding from our DAP-seq data, and protein-protein interactions (PPIs) from the STRING database ((Szklarczyk et al., 2017); **Figure 4B**).

**Figure. 4.**
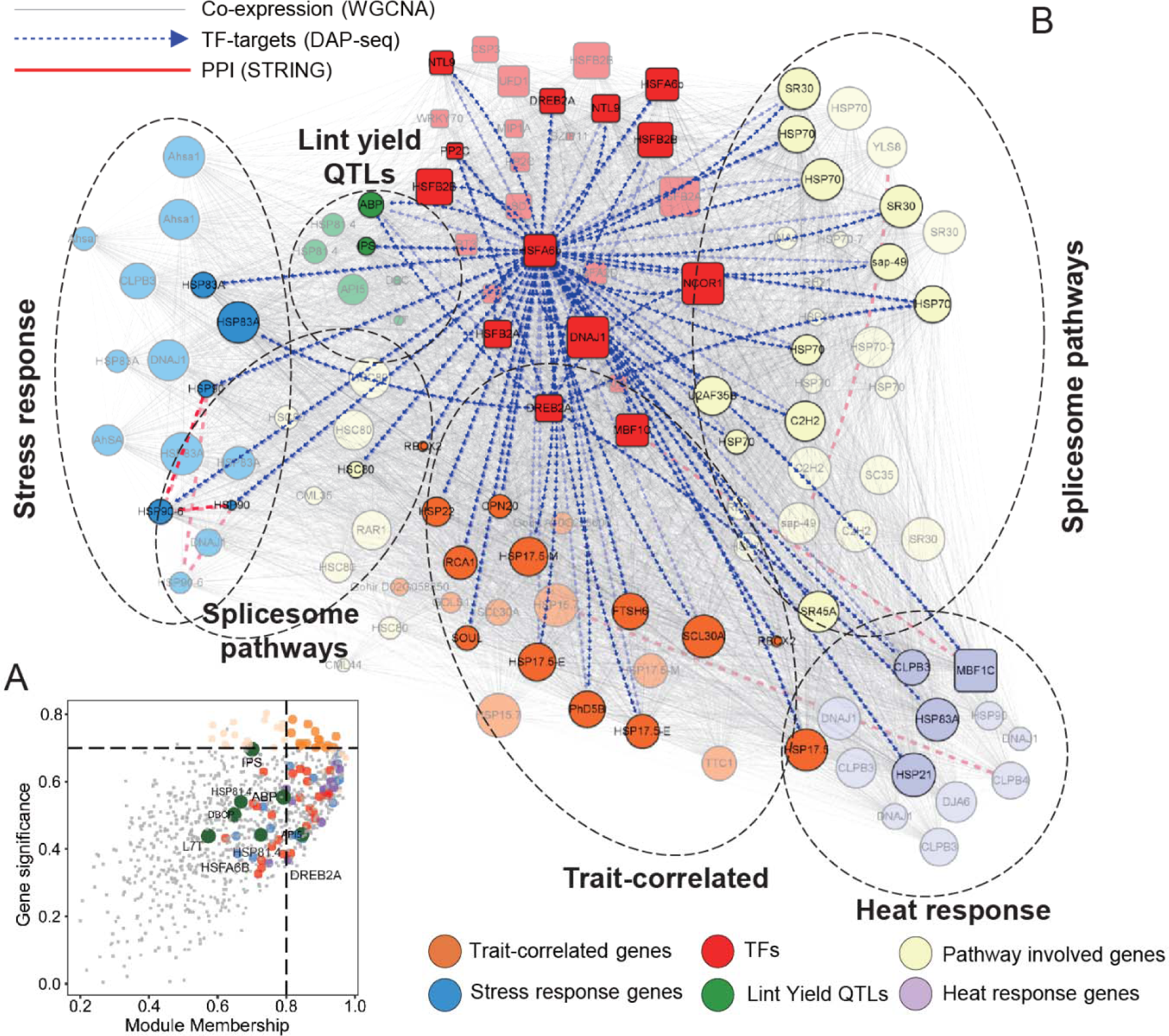
Integrated network display of a module of transcripts associated with lint yield. The module membership (MM) scores and gene significance (GS) for genes within the lint-yield module were plotted with the linear model fitted line (**A**). The dashed black lines indicate the two cutoffs used to identify trait-related genes (orange dots, *GS >* 0.7, MM > 0.8). The integrated network derived from 133 out of 866 selected genes within the lint-yield associated module were presented by layering multiple levels of information (**B**): gene categories are shown in the bottom line, as corresponded to the color of highlighted dots in MM-GS plot, including genes associated with stress response (blue), TFs (red), heat response (light purple), spliceosome pathways, biotic response pathways, and five lint yield QTLs (green), whereas connection types are shown in the top right. The solid “gray” lines connected the co-expressed genes from transcriptional level, the dashed “red” lines indicated protein-protein interactions derived from STRING database, the “blue” dash-arrowed lines highlighted targets bound by DREB2A and HSFA6B, as supported by DAP-seq data.

This network illustrates the high degree of connectivity between *GhHSFA6B-D* and the trait-correlated genes, both in terms of expression and direct connection (**Figure 4B**). Of the 24 trait-correlated genes, GhHSFA6B-D is predicted to bind 16 target genes based on DAP-seq analysis (**Figure 4B** and **Supplementary Table 7**), whereas GhDREB2A-A is predicted to only bind two genes based on AME motif enrichment. As mentioned above, GhHSFA6B-D binds to both *GhDREB2A-A* and *GhDREB2A-D* homoeologs based on DAP-seq data. In addition, GhHSFA6B-D was found to bind to its own distal promoter region, suggesting it may act in an autoregulatory loop (**Figure 4B** and **Supplementary Table 7**). Finally, despite AME predicting interactions between GhHSFA6B-D, GhDREB2A-A, and the five lint yield associated genes, only two DAP-seq derived peaks between GhHSFA6B-D and *GhIPS1-A* and *GhABP-D* were observed. As the initial AME TF predictions were made based on DAP-seq data from *Arabidopsis* (O’Malley et al., 2016), it is not unexpected that TF binding preferences will have shifted slightly for these TFs in cotton, highlighting the importance of our TF binding validation.

### DREB2A and HSFA6B affect expression of both homoleogous *ABP* loci

The inferred gene regulatory network, developed from multiple lines of evidence, suggests a direct regulatory interaction between HSFA6B, DREB2A, and two of the five lint-yield associated genes, *GhABP* and *GhIPS. GhABP-A* (Gohir.A12G006700) is an auxin-binding protein involved in cell elongation and cell division that was previously identified as a lint yield QTL (Zhu et al., 2020). TF binding motif predictions based on the Arabidopsis Cistrome Database (O’Malley et al., 2016) identified DREB2A and HSFA6B motifs for both homeologous *GhABP* loci (*GhABP-D* and *GhABP-A*; yellow boxes, **Figure S7A** and **Figure S6**). *GhABP-A*, but not *GhABP-D*, (Gohir.D12G006100; *GhABP-D*), was bound by both TFs based on our DAP-seq data (GhHSFA6B-D peak center: -561 bp TSS and GhDREB2A-A peak center: -1,445bp TSS, **Figure S7A**). *GhABP-D* was also absent from the lint yield module, and a comparison of expression versus lint yield across the 21 accessions revealed that *GhABP-D* was expressed at lower levels than *GhABP-A* (**Figure S6**). In addition, *GhABP-A* displayed a stronger correlation with the lint yield across the accessions. Despite the lower expression level of *GhABP-D* than *GhABP-A*, the abundance of both transcripts was positively correlated with both TFs (**Figure S7C** and **Figure S7**). A sequence comparison of the identified TF binding regions between *GhABP-A* and *GhABP-D* revealed only minor changes near the GhDREB2A-A (two SNPs, **Figure S7A**, top) and within the HSFA6B motifs (four SNPs, **Figure S7A**, bottom). No changes were observed within the core dehydration responsive elements (DREs) or heat shock elements (HSEs) that these two TFs are known to bind to in other systems (**Figure S7A**, red boxes).

To determine if the observed SNPs are responsible for the altered transcript abundance and DAP-seq peaks arising from these two homologous loci, we performed an electrophoretic mobility shift assay (EMSA, **Supplementary Table 11**), using ∼50-bp probes that spanned the respective DRE and HSE motifs (**Figure S7A**, pink bars). Both recombinant *GhDREB2A-A* and *GhHSFA6B-D* bound to their respective labeled probes, causing a gel shift that was not evident in the probe alone or probe + empty vector controls (**Figures S7D** and **S7E**). As expected, adding a large molar excess (200 ×) unlabeled probe to the reaction was sufficient to compete for protein binding. Interestingly, the probes corresponding to the *GhABP*-D homeolog were also able to compete for binding with GhDREB2A-A and GhHSFA6B-D (**Figures S7D** and **S7E**). These data suggest that the SNPs observed between the *GhABP*-A and *GhABP-D* distal promoter regions are insufficient to explain the specificity observed between these paralogs.

### HSFA6B affects lint yield by modulating the expression of an inositol-phosphate synthase (IPS)

The *IPS* gene, also known as MIPS in Arabidopsis, encodes for the Myo-inositol-phosphate synthase protein (INO-1), which catalyzes the rate limiting step in the synthesis of Myo-inositol-6-phosphate, a key source of phosphate in seed endosperm (Mitsuhashi et al., 2008) In addition, myo-inositol is a precursor of the osmoprotectants galactinol and raffinose, and thus is critical in a number of abiotic and biotic stress responses (Vinson et al., 2020). In contrast to ABP, there are four IPS genes in upland cotton (Gohir.D03G043600 - *GhIPS1-D*, Gohir.A02G132300 - *GhIPS1-A*, Gohir.D11G224000 - *GhIPS2-D*, and Gohir.A11G199700 - *GhIPS2-A*, **Figure S8**), two of which (*GhIPS1-D* and *GhIPS1-A*) show a positive correlation between RNA abundance and lint yield across our panel (**Figure 5B**). These two homeologous IPS genes fall in syntenic regions of the “D” and “A” subgenomes and are phylogenetically distinct from *GhIPS2-A* and *GhIPS2-D* (**Figure 6A**).

**Figure. 5.**
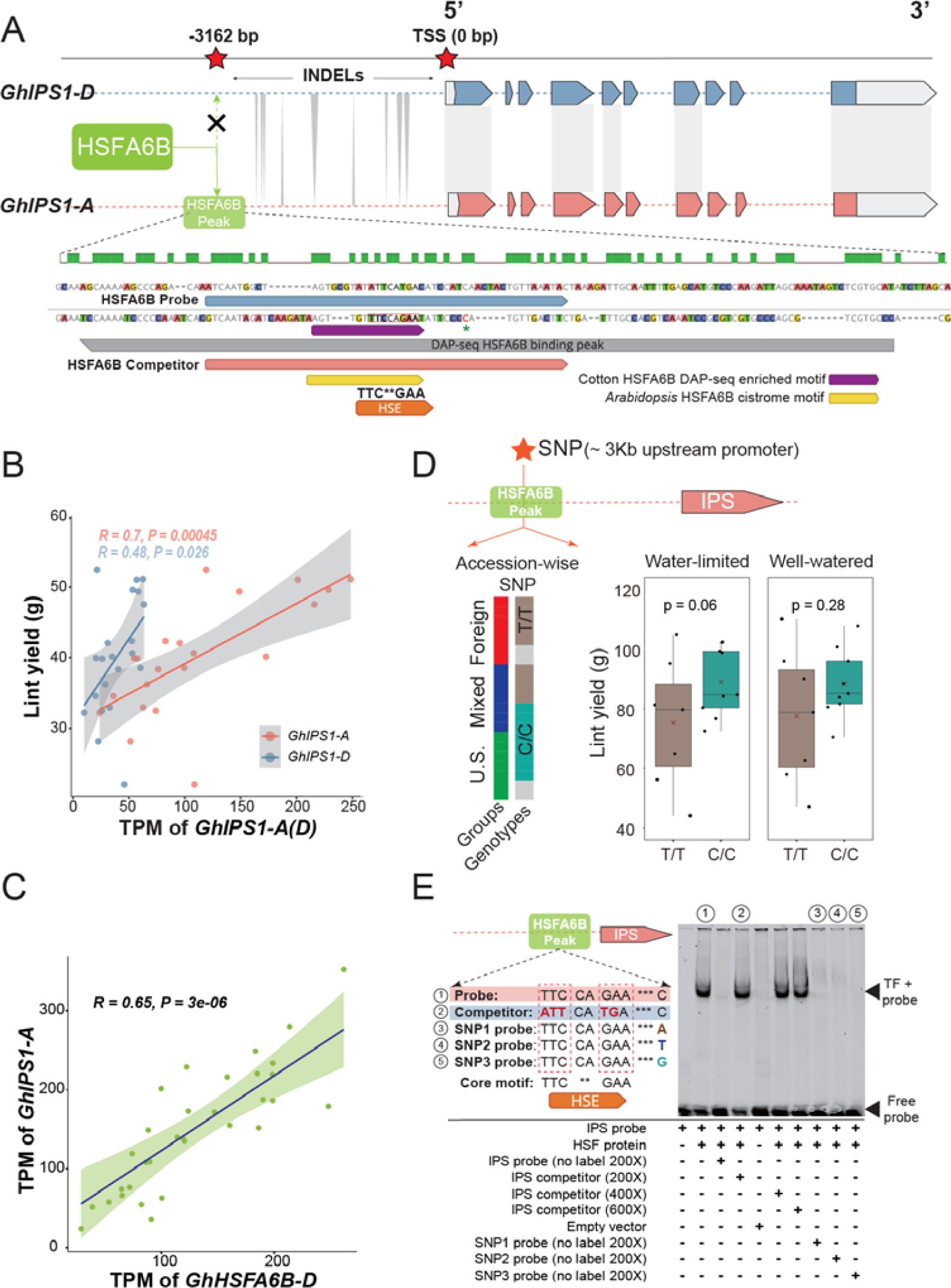
EMSA validation of interaction between *GhIPS1-A* and GhHSFA6B-D. **A) Top**: schematic representation of *GhIPS1-A* and its homeolog, *GhIPS1-D* depicting the gene structure as well as the distal promoter region where the HSFA6B binding site was identified. Filled boxes represent annotated exons. Grey shading connecting the two genes represents sequence similarity, with the distal promoter region showing high structural and sequence variation. **Bottom**: An alignment for the DAP-seq peak for HSFA6B in the promoter region of *GhIPS1-A* and its corresponding region from *GhIPS1-D*. Also shown are the DAP-seq and Cistrome identified GhHSFA6B-D binding site (HSE, purple and yellow boxes, respectively), as well as the labeled probe and competitor oligos used for EMSA. A green asterisk denotes the site of the C:T SNP observed within our panel. B) Pearson correlation between *GhIPS1-A* (red), *GhIPS1-D* (blue) expression (TPM), and lint yield. **C)** Pearson correlation between TPM values of *GhIPS1-A* and *GhHSFA6B-D*. D) Variation in lint yield between accessions associated with “C/C” (blue) and “T/T” (brown) genotypes of a SNP adjacent to the HSE element in the distal promoter of *GhIPS1-A* (∼3.1 Kb upstream of *GhIPS1-A* start). Accessions lacking sufficient coverage at this site are shown in grey. Accessions bearing the C/C or T/T genotypes were also divided based on their phylogenetic groupings (i.e., “U.S.”, “Foreign”, and “Mixed”) from **Figure 1A**. E) Interaction between the GhHSFA6B-A protein and *GhIPS1-A* probe was examined by EMSA. The reaction components for each lane are listed below the gel image, including the IRD700 labeled probe, unlabeled probe (200×, same sequence as the labeled probe), competitor sequence (200×, 400×, and 600×, containing the *GhIPS1-D* disrupted HSE), empty vector, and the other competitor sequence (200×, containing one of the three SNP probes, A, T, or G).

**Figure. 6.**
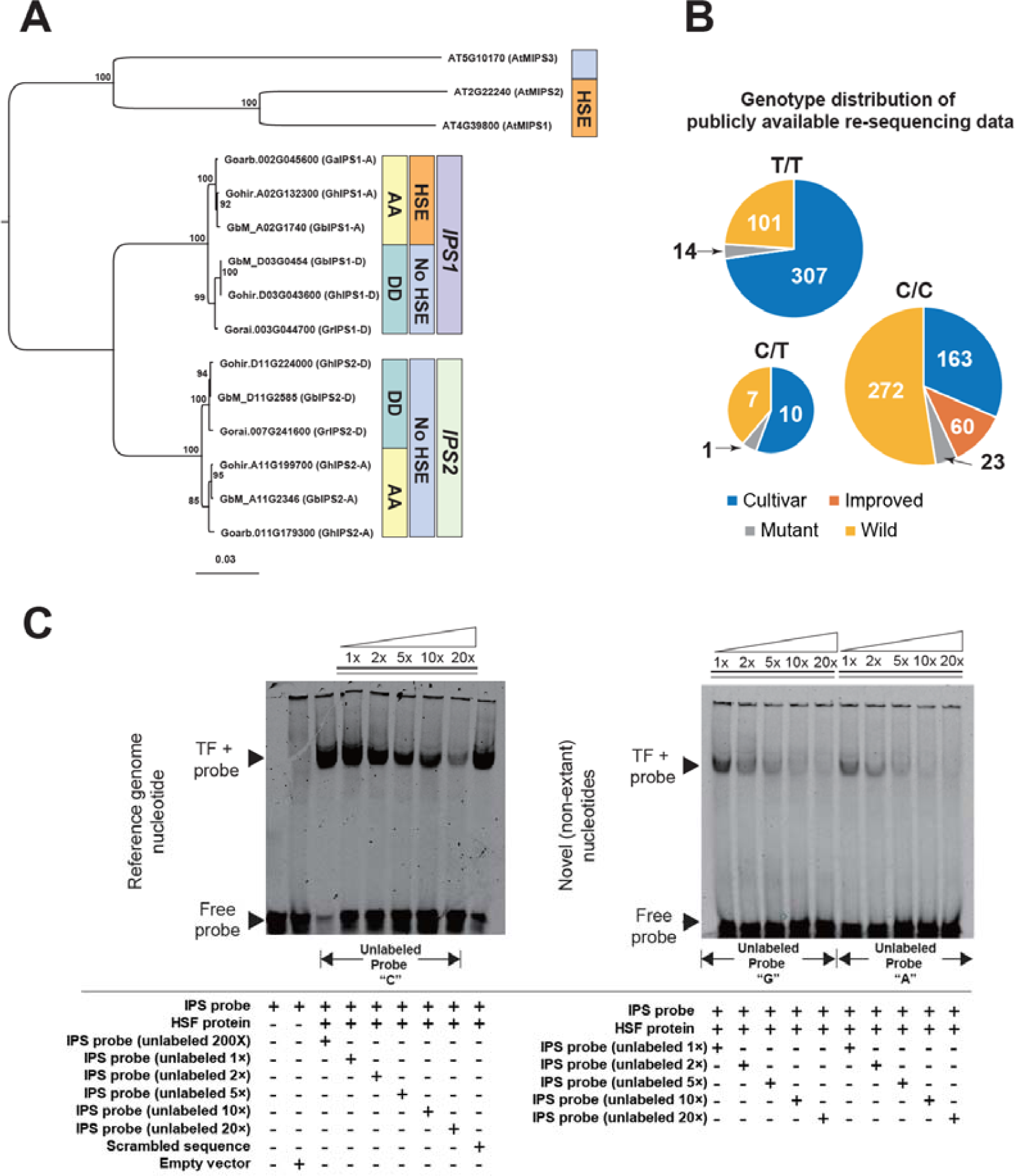
Identification of a HSE-adjacent SNP that impacts GhHSFA6B-D binding to GhIPS1-A. **A)** A phylogenetic tree containing *GhIPS1* and *GhIPS2* homeologous genes from *G. hirsutum* (AADD), *G. barbadense* (AADD), *G. arboreum* (AA), and *G. raimondii* (DD) was constructed using *Arabidopsis thaliana* IPS homologs as an outgroup. The “orange” and “light blue” label denotes the presence of the HSE upstream of *GhIPS1-A* genes in all AA genome containing species, but not in the distal promoters of *GhIPS1-D* genes found in DD genome species, nor in any of the the *GhIPS2* paralogs in either AA or DD genomes. **B**) Distribution of genotypes of the *GhIPS1-A* DAP-seq peak-associated SNP (C:T) in a published whole genome sequencing-based panel (**1024 accessions Yuan et al., 2021**). The distribution is summarized in pie-charts reflecting the accession frequencies of the wild, cultivated, mutants, and improved *G. hirsutum* accessions across the “**C/C**” genotype (reference genome genotype), “**T/T**” genotype, and “**T/C**” genotype. **C**) The level of the GhHSFA6B-D to *GhIPS1-A* binding efficiency of different IPS-associated SNP variants was examined by competitions between the IDR700 dye-labeled reference oligos and non-labeled oligos with the reference nucleotide “C”, or the non-native SNP oligos “G”, and “A”, respectively. Left panel: binding efficiency of the reference “C” containing probe was tested by titrating in increasing amounts of the competitor probe (unlabeled reference “C”). **Right panel**: the impact of this nucleotide position on GhHSFA6B-D binding was tested by competing the labeled reference probe with increasing amounts of two non-native probes (“G” or “A”). Components in each reaction are shown below.

Despite a reported expression bias towards D subgenome homeologs (Chen et al., 2020), only *GhIPS1-A* was predicted to contain a GhHSFA6B-D binding site based on DAP-seq data and the Arabidopsis Cistrome Database (**Figure 5A**, purple and yellow boxes, respectively). A pairwise comparison of the distal promoter regions of *GhIPS1-A* and *GhIPS1-D* revealed substantial polymorphisms between the two regions that appear to have disrupted the core HSE in this region in *GhIPS1*-*D*, as well as between *GhIPS1*-*D* and *GhIPS2*-*A/D* (**Figure 5A**, bottom; **Figure S8A**). An examination of *IPS1* distal promoter elements in *G. hirsutum, G. barbadensis, G. raimondii (*a D-subgenome representative) and G. arboreum (old-world cotton and an A-subgenome representative) revealed conservation of the HSE within *GhIPS1*-*A* in *G. hirsutum, G. barbadensis*, and *G. arboreum*, but not in *IPS1-D* loci for any of these species (**Figure 6A**). In agreement with the DAP-seq data, we observe a strong positive correlation between *GhHSFA6B-D* and *GhIPS1-A* transcript abundance, but not between *GhHSFA6B-D* and any of the other *GhIPS* loci (**Figure 5C; Figure S7**). Interestingly, an HSE, and AtHSFA6B DAP-seq peak, were observed upstream of the Arabidopsis IPS1 and IPS2 paralogs in Arabidopsis Cistrome data (At4G39800 and At2G22240, respectively; **Figure 6A; Figure S10**), suggesting this regulatory mechanism may be conserved between these two species.

An examination of polymorphisms within our RNA-seq data relative to the *TM-1* reference genome uncovered a lint-yield associated SNP directly adjacent to the predicted GhHSFA6B-A binding motif (**Figure 5A**). While this nucleotide is a cytosine (C) at this position in the reference genome and in *G. barbadensis* and *G. arboreum*, we observed two genotypes in our panel, either with a cytosine (C, *n* = 8) or a thymine (T, *n* = 9; **Figure 5D**). Interestingly, the “C” genotypes, which were either the US or “mixed” accessions, showed increased lint yield under water-limited conditions relative to the “T” genotypes (**Figure 5D**), suggesting this region might impact GhHSFA6B-D binding. To define the GhHSFA6B-D binding site more carefully in the *GhIPS1-A* promoter region, we designed a labeled probe centered on the core HSE (**Figure 5A**, pink box). A gel shift was observed when this probe was combined with *in-vitro* expressed GhHSFA6B-D protein (**Figure 5E**). As expected, this signal was abolished when a large excess of unlabeled probe was added. The addition of an unlabeled competitor, corresponding to the homologous *GhIPS1*-*A* promoter region with a disrupted HSE (Oligo 2, **Figure 5E**) was unable to abolish binding, even when adding a large molar excess (400**×** and 600**×**; **Figure 5E**). As the labeled probe contained the reference nucleotide (C) at the site of the observed lint yield-associated SNP, we next tested if altering this nucleotide, but leaving the rest of the HSE intact, had an impact on HSFA6B binding. Oligos with this site altered to an A, T, or G nucleotide (SNP probes 1-3) could compete for GhHSFA6B-D binding when added in excess (200**×**; Figure 5E).

While the SNP oligos were competing with the reference oligo when added in excess, due to its location outside of the HSE it is not clear if the site of the SNP impacts GhHSFA6B-D binding. To precisely address this question, we performed a competition experiment using lower concentrations of two non-native SNP probes, SNP-A and SNP-G (SNP probes 1 and 3), that are not present in the genomes of our panel or a much larger sequenced panel comprised of 1024 accessions (Yuan et al., 2021)**, Supplementary Table 12**). Surprisingly, equimolar amounts of either non-native competitor SNP oligos were capable of competing with the reference probe to a higher degree than the unlabeled reference oligo (**Figure 5B**, **Figure 6C, and Figure S9**). These data suggest that this site, which contains the only observed SNP within the DAP-seq peak region of *GhIPS1*-*A* in our panel, has a strong influence on GhHSFA6B protein binding, *GhIPS1*-*A* expression, and lint yield. Given the differentially accumulated frequency of SNPs at this site in extant accessions (wild, cultivated, improved, and mutant) (**Figure 6B**), this site has likely been under different levels of selection in wild and cultivated accessions due to its connection to improved yield.

### Conclusions

Understanding the regulatory crosstalk between agronomic traits of interest (e.g., yield) and heat and drought stress responses are critical for developing drought-tolerant cultivars with minimal impacts on yield (Alizadeh et al., 2020). Characterizing genotype-specific molecular responses under water-limiting conditions is challenging due to the genetic complexity underlying quantitative physiological traits (Welcker et al., 2011; Tardieu et al., 2011). In most crop species this genetic complexity is influenced at the pre- and post-transcriptional level by *cis*-regulatory control, gene structural variation, and post-translational modifications (Tardieu et al., 2017; Joshi et al., 2016). In cotton, this complexity is further compounded by a recent (∼1.0-1.6 Ma) allopolyploidy and strong human selection within the last 8,000 years (Chen et al., 2020). Despite the strong selection existing in current elite *G. hirsutum* cultivars relative to other domesticated crops (Chen et al., 2020; Ma et al., 2018; Su et al., 2016a), variation in gene expression was identified across the panel for a cohort of transcripts that could be associated with improved yield under water limiting conditions.

Despite the developmental stage at which we sampled for transcriptomics (primarily the cell elongation stage within the developing bolls; Ma et al., 2018) there was already a strong transcriptomic signal in our data connecting abiotic stress factors and lint yield. This group of lint yield and abiotic stress-associated genes were largely regulated by the well-known transcription factors *GhDREB2A-A* and *GhHSFA6B-D*. Importantly, based on DAP-seq, motif enrichment, and co-expression, HSFA6B not only regulates *GhDREB2A-A* in this module, but also regulates itself and two lint yield-associated genes, *GhABP* and *GhIPS1-A*. While *GhDREB2A-A* was previously shown to be regulated by *GhHSFA6B-D* in *Arabidopsis*, this regulation was dependent on ABA in general, and specifically the ABA response element TF AREB1 (Huang et al., 2016; Sakuma et al., 2006). In contrast, in cotton this regulation does not appear to be ABA-dependent, as none of the typical ABA-responsive elements, nor ABA biosynthesis genes, were associated with this module, nor does HSFA6B contain a canonical AREB binding domain (ABRE) or share an expression profile with GhAREB1. However, this hypothesis requires further examination. This specialized regulatory mechanism appears to have emerged in cotton well before domestication. Indeed, the old-world variant of IPS1 arising from the A subgenome (*GhIPS1-A*) is the only IPS to be targeted by *GhHSFA6B-D*. In fact, this HSFA6B – *IPS1* regulatory connection may reflect an evolutionarily conserved stress response mechanism, as it is also observed in Arabidopsis. Of importance to breeders, this regulatory mechanism appears to have undergone additional selection since speciation to maintain yield under water limited conditions.

## Author Contributions and Acknowledgements

LY, ADLN, and DP developed the project. AND performed RNA extractions and library prep, GM performed metabolomics analysis. KRT contributed to field data collection. XZ performed the DAP-seq library prep. LY, VPTK and XZ conducted the EMSA experiments. KRT and LH contributed to the execution of the field trial. LH contributed to the germplasm. LY, GM, DP, and ADLN contributed to the writing of the manuscript. All authors contributed to the review and approval of the manuscript. The authors would like to acknowledge support from the NSF-IOS #2023310 (awarded to ADLN), NSF-IOS #2102120 (awarded to DP and ADLN), Cotton Incorporated Core Projects 18-384, 20-720, and 23-890 (to DP) and 22-639 (to ADLN), and the United States Department of Agriculture-Agricultural Research Service (USDA-ARS) (KT, CRIS #2020-13660-008-00-D; and LH, CRIS #3091-21000-041-00D). Mention of trade names or commercial products in this publication was solely for the purpose of providing specific information and does not imply recommendation or endorsement by the USDA. The USDA was an equal opportunity provider and employer. The authors would like to thank other members of the Nelson, Skirycz, and Pauli labs, as well as Dr. Jeffrey Chen (UT-Austin) for helpful discussion in the formulation of the analyses presented herein. The authors declare no competing interests.

## Data availability

The Illumina reads of transcriptome data have been deposited in the NCBI database under BioProject submission number SUB11686475. Analysis code is available here: https://github.com/Leon-Yu0320/Cotton_omics_studies.

## Supplementary Figures

**Figure. S1 Comparison of six production-related traits and four vegetative indices from two watering conditions**

The 21 upland cotton accessions were split into “Foreign”, “U.S.”, and “Mixed” groups for pairwise comparisons among 10 traits from two water conditions. For each pair-wise comparison, a t-test was performed to effects of combined heat-drought stress.

**Figure. S2 Transcriptome comparison of expression profiles in the three phylogenetic groups**

**A)** Upset plot of differentially expressed (DEGs) in all representatives within a particular phylogenetic group. The vertical black bars stand for the numbers of unique DEGs from each group, intersection between each two group, and intersection among three groups.

**B)** Enriched gene ontology terms of commonly down-regulated (top) and up-regulated (bottom) genes among 21 accessions. The calculation is based on -log10 Q-values which represents the level of enrichment (value cutoff > 3).

**Figure. S3 Weighted Co-expression network and trait-module membership**

**A)** The left panel represents the frequency of gene connection revealed by connectivity (k). The right panel shows the log-log plot of the connectivity, the correlation coefficient represents the scale of free topology. The 0.86 here indicated the high approximation of a scale-free of network of network derived from top 10% MAD score.

**B)** The left panel is the relation between approximation of a scale-free of network and different soft threshold (power). The seven is the least numbers to ensure a scale free network. The right panel stands for the mean connectivity of network when different power score was applied.

**C)** A heatmap plot depicting the topological overlap matrix (TOM) supplemented by hierarchical clustering dendrograms and the module colors was created by TOMplot function from the WGCNA package. Each cell corresponds to a single gene, the light color indicated the high topological overlap, and the darker orange and red colors represent the lower topological overlap.

**Figure. S4 Preparation of DAP-seq libraries**

**A)** The genomic DNA of the TM-1 cotton genome was extracted with the gel running before and after the adaptor ligation;

**B)** The immunoblots of the HSFA6B and DREB2A confirms the expected molecular mass; The protein purification efficiency was monitored at all steps from antibody coupled beads binding, washing and elution.Examination of the efficacy of upon antibody coupled beads binding, washing, and protein purification.

**Figure. S5 EMSA validation of interaction between ABP and two TFs**

Sequence alignment of *GhABP* and an *GhABP* paralog: promoter and coding region of a lint yield candidate ABP and its paralog were aligned using MUSCLE (iteration = 200), two peaks bound by *GhDREB2A-A* and *GhHSFA6B-D* were identified around the 1.5 Kb upstream of promoter regions (marked by red pentagram). Annotation of the peak region revealed three tandem heat shock elements (HSE, orange bracket), which overlapped with the motif region detected by FIMO and MEME (purple and yellow bracket respectively). SNPs were detected within one HSE (orange box) while the rest HSEs are identical from two paralogs. One EMSA DNA probe and competitor pair was designed to span two identical HSEs and the other probe-competitor pair to cover the HSE bearing SNPs (bottom panel). Similarly, the *GhDREB2A-A* peak region was annotated and marked (top panel).

**A)** Pearson correlation between TPM values of *GhABP-A* and its two putative regulating TFs, *GhDREB2A-A* and *GhHSFA6B-D*.

**B)** Pearson correlation between the expression (TPM) *GhABP-A* (red) and its homeolog, *GhABP-D* (blue), with lint yield.

**C)** Interaction between *GhDREB2A-A* and *GhABP-A* was examined by EMSA. The reaction components for each lane are listed below, including the IRD700 labeled probe, unlabeled probe (200×), competitor sequences (200×, containing 2 SNPs flanking the HSE), and empty vector.

**D)** Interaction between *GhHSFA6B-D* and *GhABP-A* was examined by EMSA. The reaction components for each lane are listed below, including the IRD700 labeled probe, unlabeled probe (200×), competitor sequences (200×, containing 3 SNPs flanking the HSE), and empty vector.

**Figure. S6 Distribution of GhDREB2A and GhHSFA6B DAPseq peaks**

An IGV screen shot revealed the binding region (peak) across the distal promoter regions of ABP (bound by both *GhDREB2A-A* and *GhHSFA6B-D*), IPS (bound by *GhHSFA6B-D*), and *GhDREB2A-A* (bound by *GhHSFA6B-D*) across two replicates of DAP-seq libraries.

**Figure. S7 Correlation between TFs and target genes (TPM)**

Pearson correlation between TPM values of *GhIPS-A (D)* genes and HSFA6B, TPM values of *GhABP-A (D)* genes and *GhHSFA6B-D*, and TPM values of *GhABP-A (D)* genes and *GhDREB2A-A*.

**Figure. S8 Sequence alignment and expression profile of four IPS in cotton**

**A)** Sequence alignment of Four IPSs identified in upland cotton genome: promoter (left panel) and coding region (right panel) were aligned using MUSCLE (iteration = 200), only one peak bound by *GhHSFA6B-D* was identified around the 3.1 Kb upstream of promoter regions (marked by red pentagram).

**B)** Correlation between TPM value of four *GhIPSs* and lint yield, tested by Pearson correlation among 21 accessions.

**Figure. S9 Comparison of HSFA6B - IPS binding among three genotypes**

The band area of “Protein + probe” and “probe” region were extracted by calculating the pixels of bands using ImageJ software. To illustrate the variations among three types of nucleotides (reference genotype “C” and two naturally non-existed genotypes “G” and “A”) across a concentration gradient of non-labeled probe (1**×**, 2**×**, 5**×**, 10**×**, 20**×** relative to labeled probe, **X-axis in the line graph)**, we used the ratio = (probe + protein) / probe (**Y-axis in the line graph**) to represent the level of competition between labeled probe and non-labeled probe. The higher ratio indicates the lower chances of the binding of non-labeled probe.

**Figure. S10 Screenshot of AtHSFA6B DAP-seq peaks in AtIPS paralogs.**

10:1 zoom level screenshot from neomorph.salk.edu for all HSFA6B DAP-seq binding data for AtIPS1 (**A**, AT4G39800), AtIPS2 (**B**, AT2G22240), and AtIPS3 (**C**, AT5G10170). Where distances aren’t clear, the exact coordinates of the HSFA6B and transcription start site (e.g., AT4G39800) the distances are annotated.

