## Supplemental figures for "Regulation of a single Inositol 1-Phosphate Synthase homeolog by HSFA6B contributes to fiber yield maintenance under drought conditions in upland cotton"

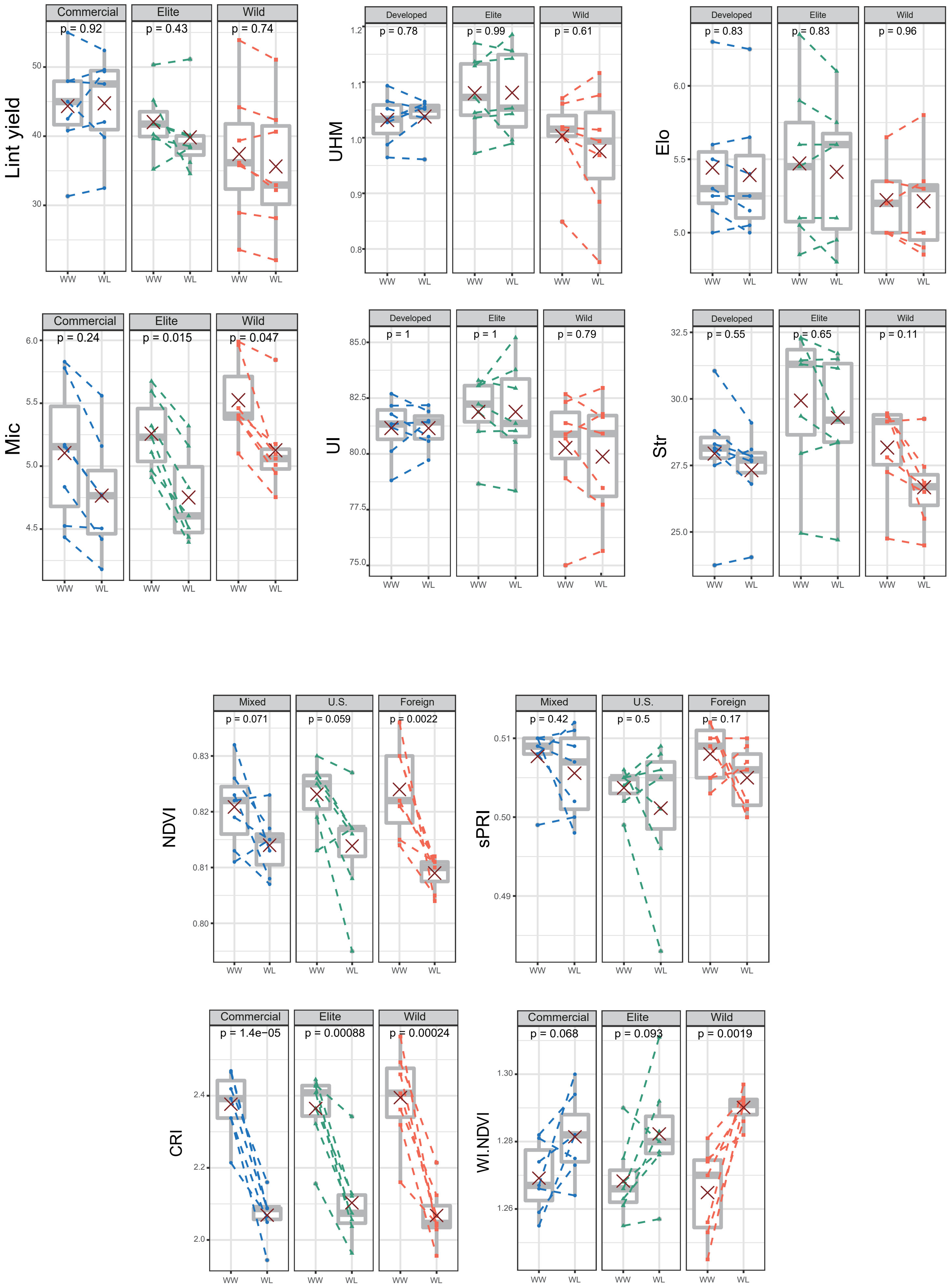
Lint yield
Commercial
p = 0.92
Elite
p = 0.43
Wild
p = 0.74
UHM
Developed
p = 0.78
Elite
p = 0.99
Wild
p = 0.61
Elo
Developed
p = 0.83
Elite
p = 0.83
Wild
p = 0.96
Mic
Commercial
p = 0.24
Elite
p = 0.015
Wild
p = 0.047
UI
Developed
p = 1
Elite
p = 1
Wild
p = 0.79
Str
Developed
p = 0.55
Elite
p = 0.65
Wild
p = 0.11
NDVI
Mixed
p = 0.071
U.S.
p = 0.059
Foreign
p = 0.0022
sPRI
Mixed
p = 0.42
U.S.
p = 0.5
Foreign
p = 0.17
CRI
Commercial
p = 1.4e−05
Elite
p = 0.00088
Wild
p = 0.00024
WI.NDVI
Commercial
p = 0.068
Elite
p = 0.093
Wild
p = 0.0019
WW
WL

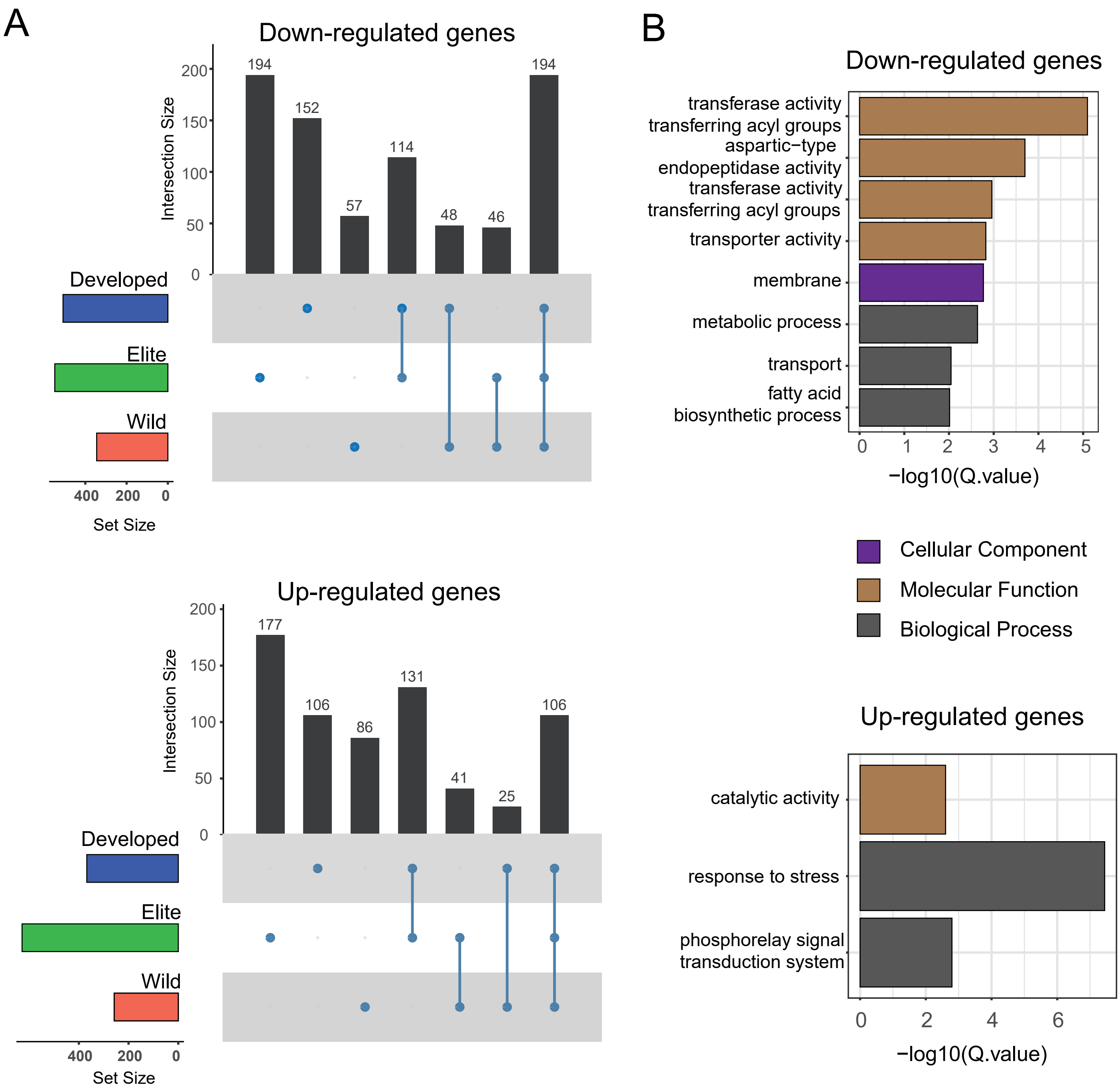
A
Down-regulated genes
Intersection Size
194
152
57
114
48
46
194
Developed
Elite
Wild
Set Size
Up-regulated genes
Intersection Size
177
106
86
131
41
25
106
Developed
Elite
Wild
Set Size
B
Down-regulated genes
transferase activity transferring acyl groups
aspartic−type endopeptidase activity
transferase activity transferring acyl groups
transporter activity
membrane
metabolic process
transport
fatty acid biosynthetic process
−log10(Q.value)
Cellular Component
Molecular Function
Biological Process
Up-regulated genes
catalytic activity
response to stress
phosphorelay signal transduction system
−log10(Q.value)

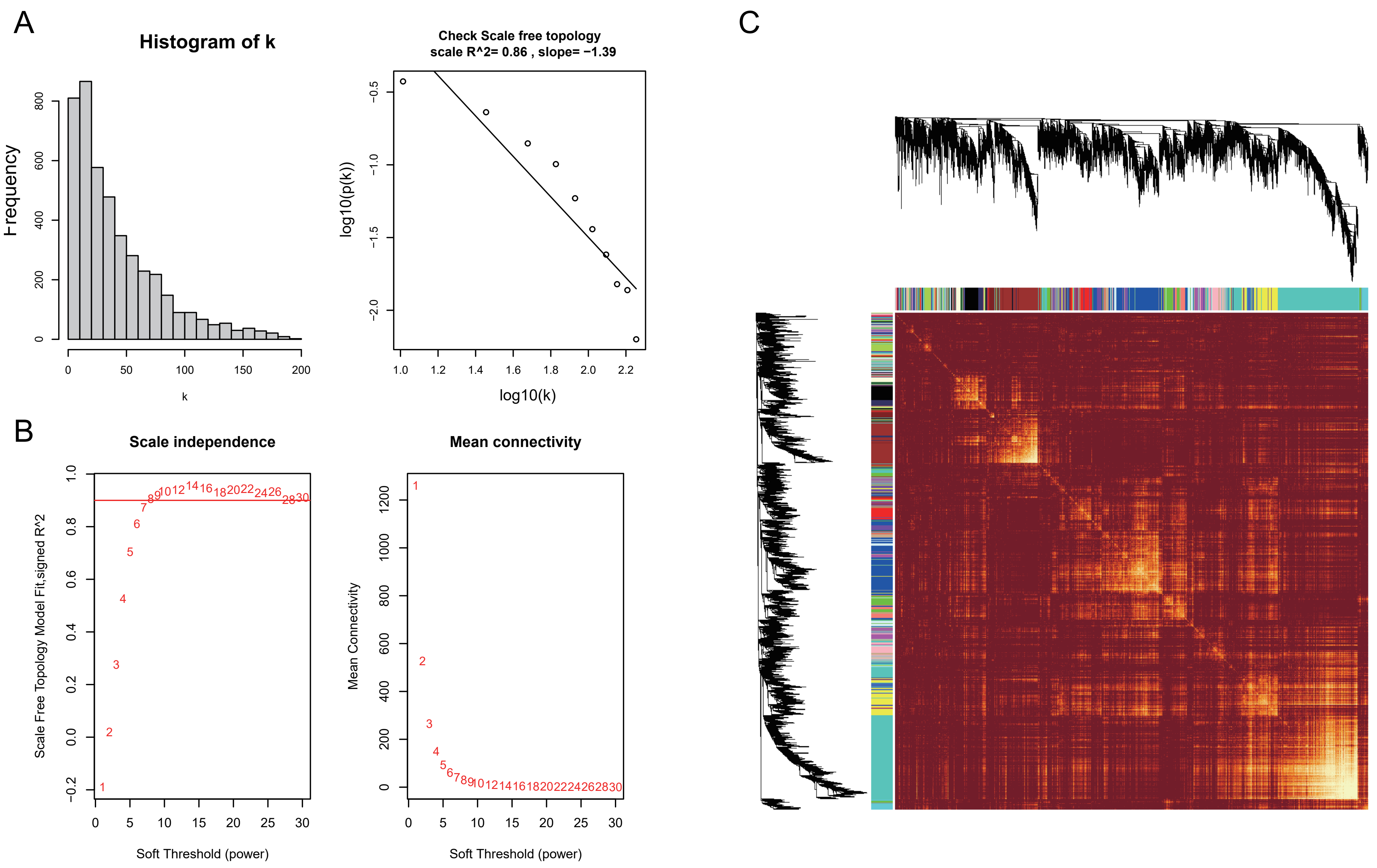
A
Histogram of k
Frequency
k
Check Scale free topology
scale R^2= 0.86 , slope= −1.39
log10(p(k))
log10(k)
B
Scale independence
Scale Free Topology Model Fit,signed R^2
Soft Threshold (power)
Mean connectivity
Mean Connectivity
Soft Threshold (power)
C

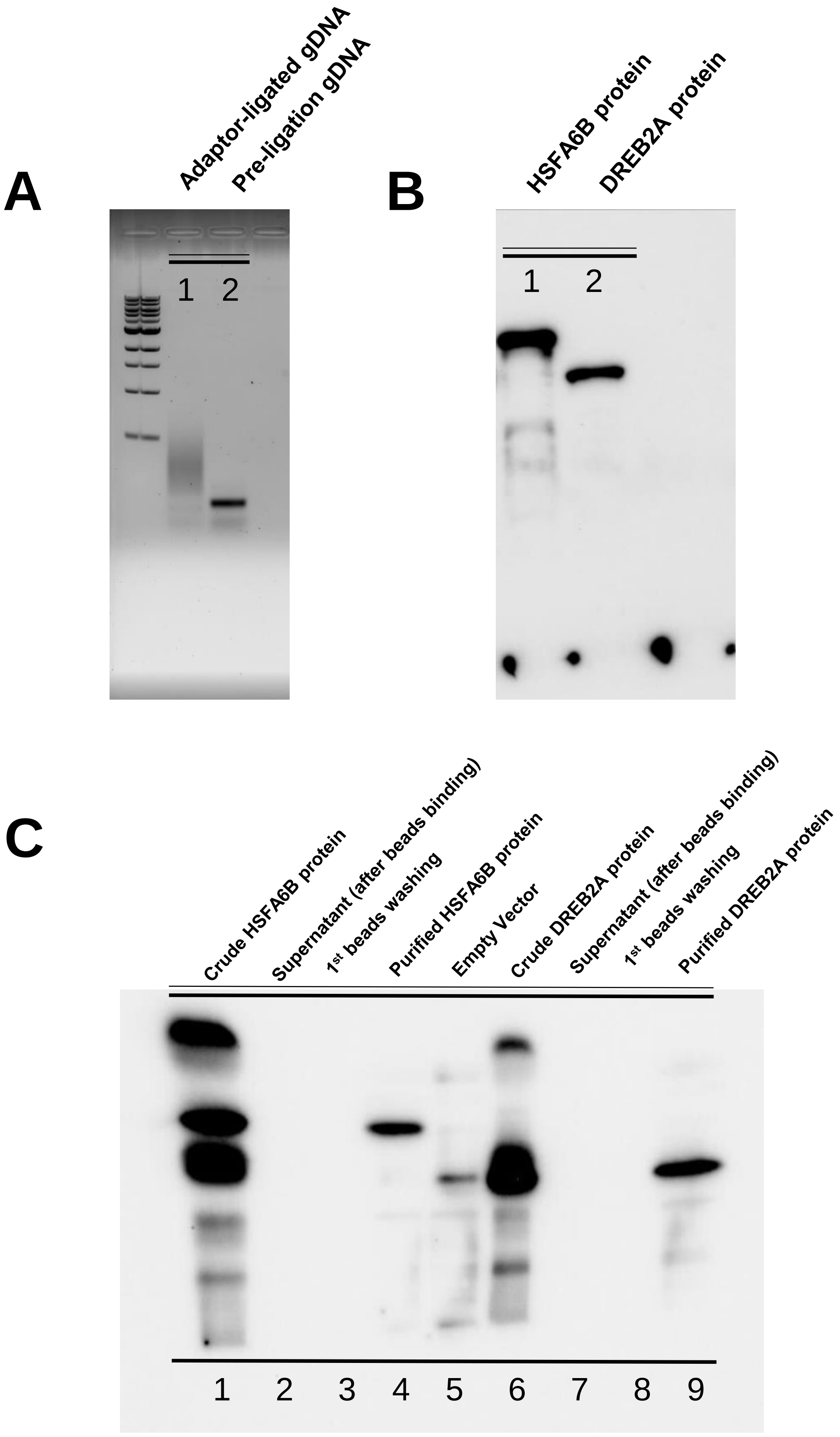
A
Adaptor-ligated gDNA
Pre-ligation gDNA
1 2
B
HSFA6B protein
DREB2A protein
1 2
C
Crude HSFA6B protein
Supernatant (after beads binding)
1st beads washing
Purified HSFA6B protein
Empty Vector
Crude DREB2A protein
Supernatant (after beads binding)
1st beads washing
Purified DREB2A protein
1 2 3 4 5 6 7 8 9

### DREB2A (A-genome) targeted by HSFA6B

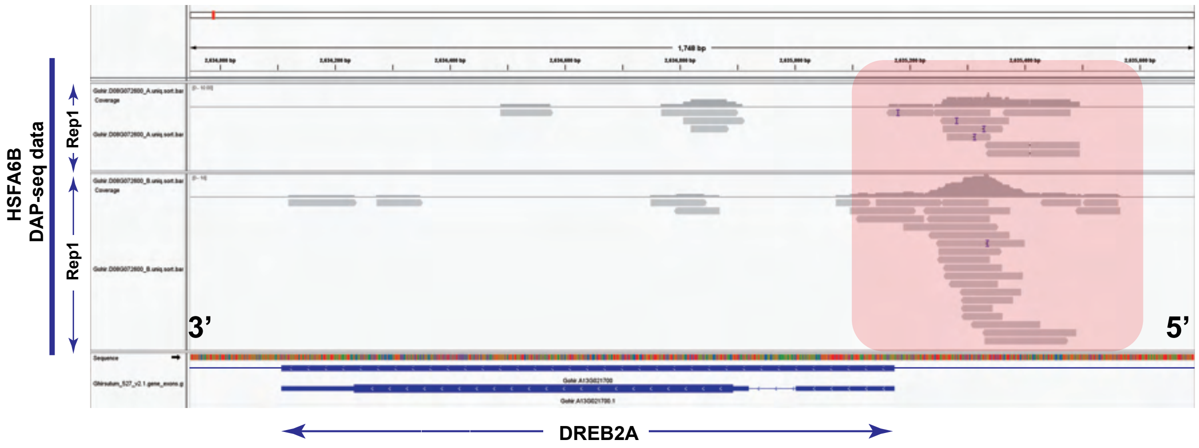


### DREB2A (D-genome) targeted by HSFA6B

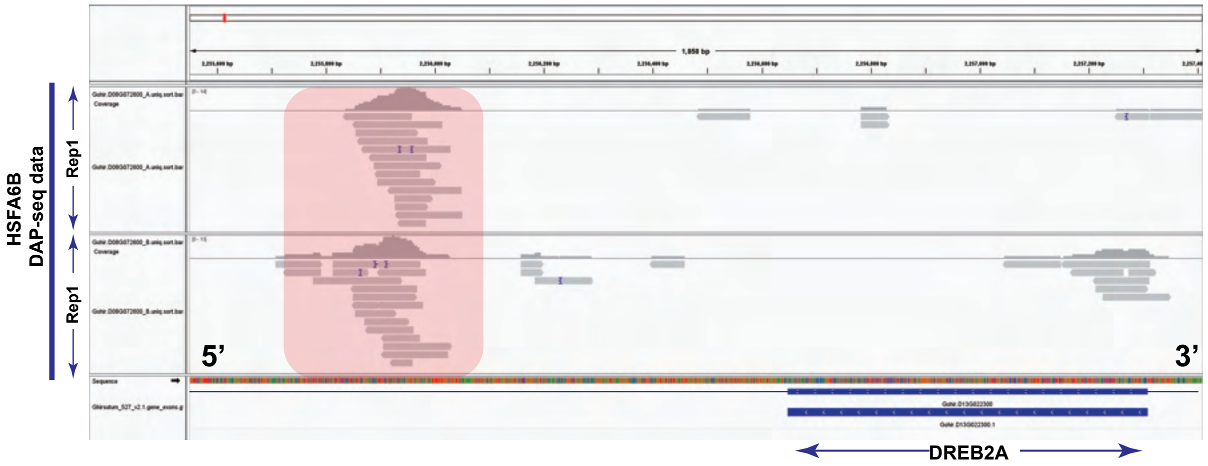


### IPS targeted by HSFA6B

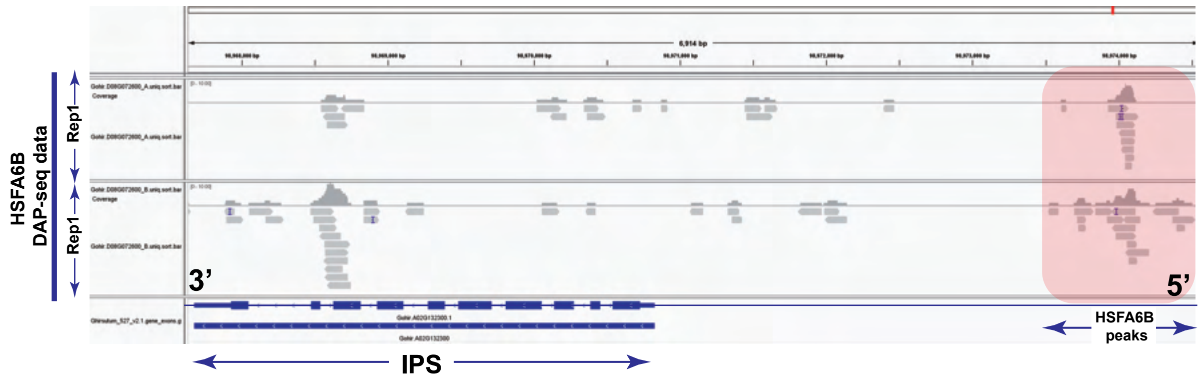


### ABP targeted by HSFA6B and DREB2A

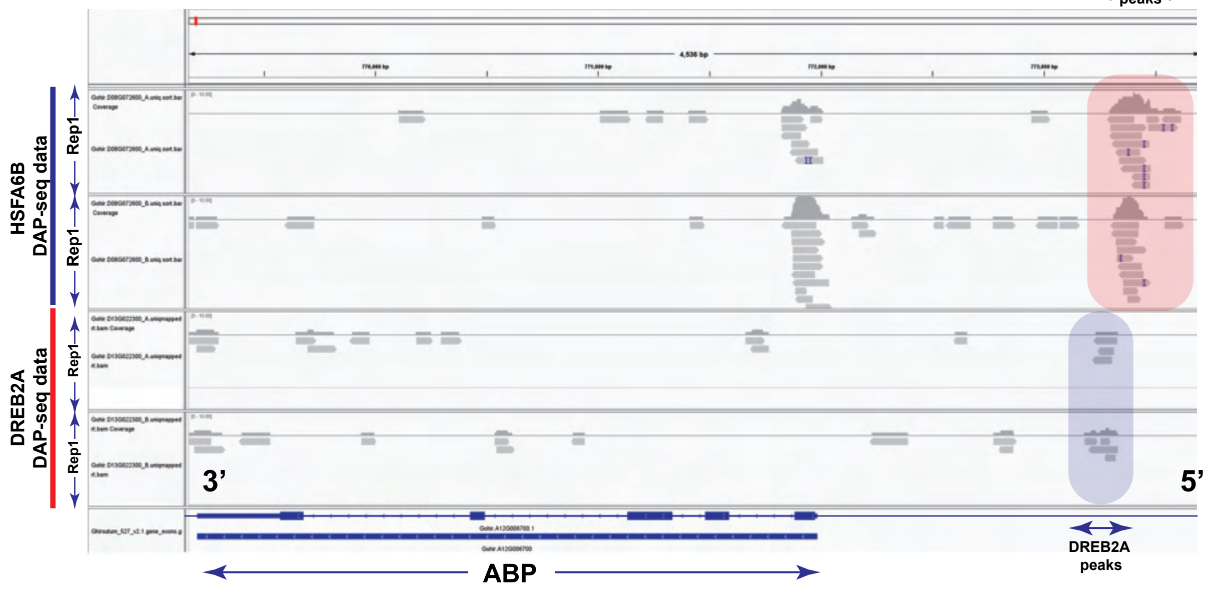

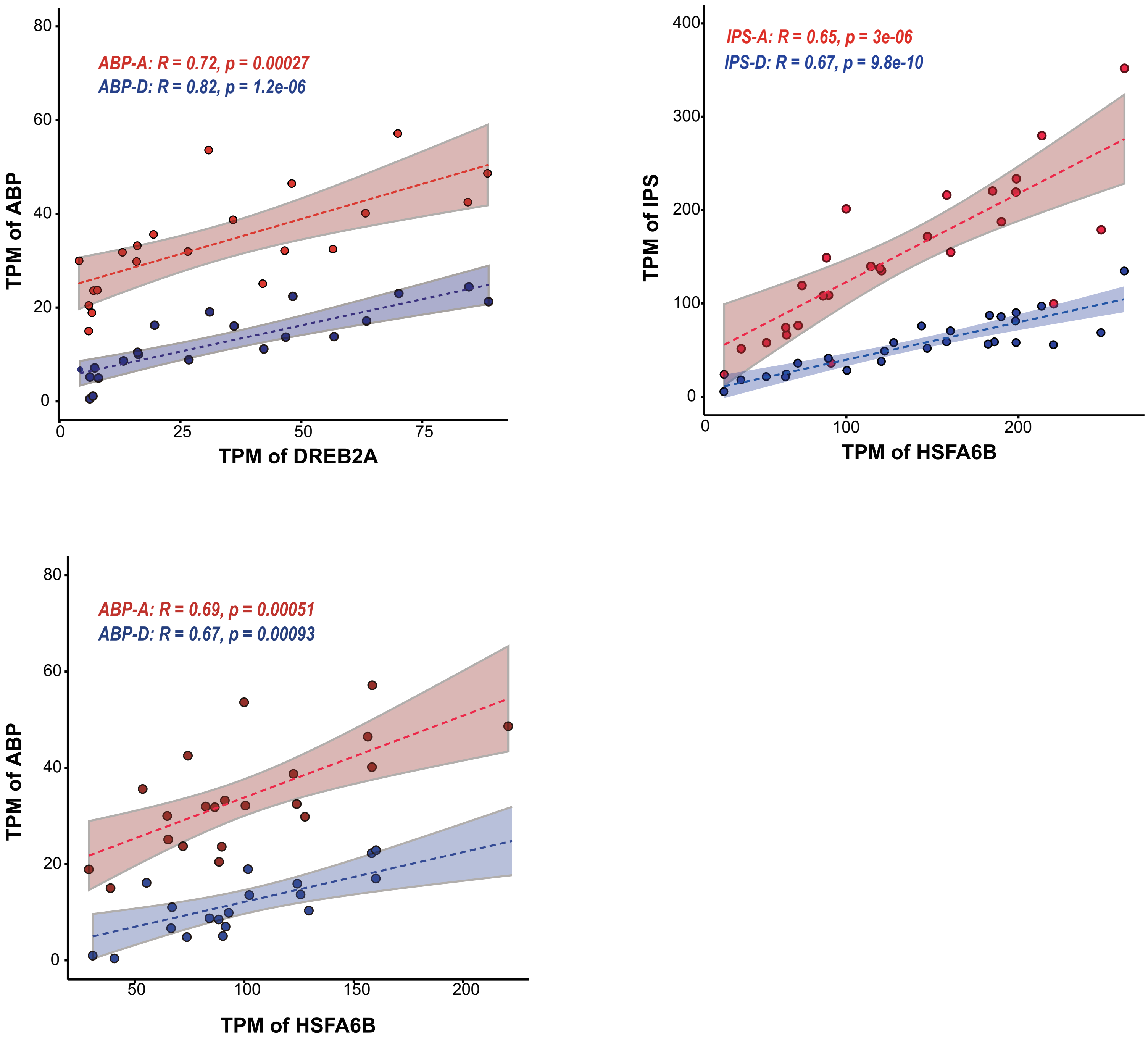

ABP-A: R = 0.72, p = 0.00027
ABP-D: R = 0.82, p = 1.2e-06
TPM of ABP
TPM of DREB2A
IPS-A: R = 0.65, p = 3e-06
IPS-D: R = 0.67, p = 9.8e-10
TPM of IPS
TPM of HSFA6B
ABP-A: R = 0.69, p = 0.00051
ABP-D: R = 0.67, p = 0.00093
TPM of ABP
TPM of HSFA6B

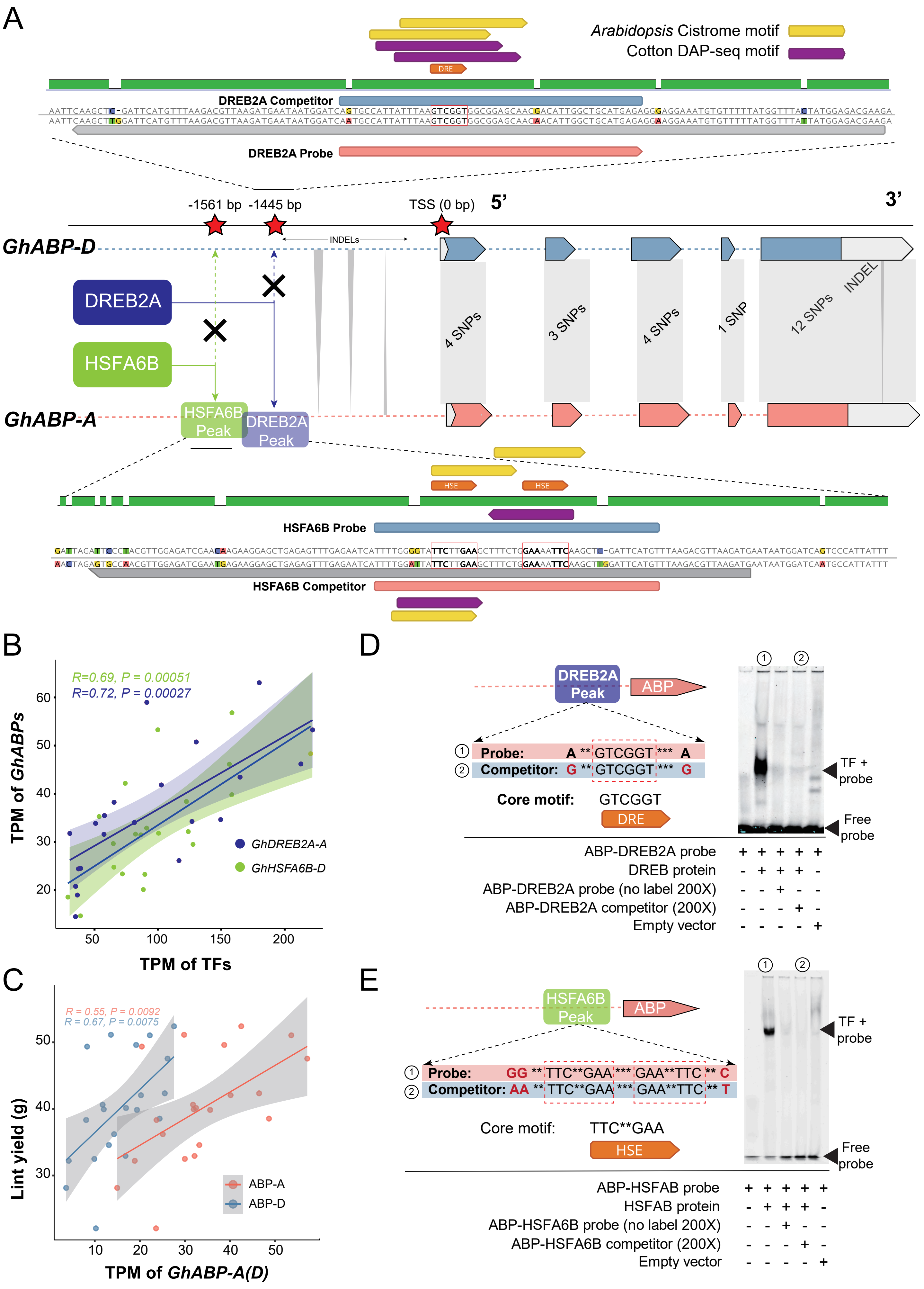
A
Arabidopsis Cistrome motif
Cotton DAP-seq motif
DRE
DREB2A Competitor
DREB2A Probe
-1561 bp
-1445 bp
TSS (0 bp)
5'
3'
GhABP-D
INDELs
DREB2A
HSFA6B
4 SNPs
3 SNPs
4 SNPs
1 SNP
12 SNPs
INDEL
GhABP-A
HSFA6B Peak
DREB2A Peak
HSE
HSE
HSFA6B Probe
HSFA6B Competitor
B
R=0.69, P = 0.00051
R=0.72, P = 0.00027
TPM of GhABPs
GhDREB2A-A
GhHSFA6B-D
TPM of TFs
C
R = 0.55, P = 0.0092
R = 0.67, P = 0.0075
Lint yield (g)
ABP-A
ABP-D
TPM of GhABP-A(D)
D
DREB2A Peak
ABP
① Probe: A ** GTCGGT *** A
② Competitor: G ** GTCGGT *** G
Core motif: GTCGGT
DRE
TF + probe
Free probe
ABP-DREB2A probe + + + + +
DREB protein - + + + -
ABP-DREB2A probe (no label 200X) - - + - -
ABP-DREB2A competitor (200X) - - - + -
Empty vector - - - - +
E
HSFA6B Peak
ABP
① Probe: GG ** TTC**GAA *** GAA**TTC ** C
② Competitor: AA ** TTC**GAA *** GAA**TTC ** T
Core motif: TTC**GAA
HSE
TF + probe
Free probe
ABP-HSFAB probe + + + + +
HSFAB protein - + + + -
ABP-HSFA6B probe (no label 200X) - - + - -
ABP-HSFA6B competitor (200X) - - - + -
Empty vector - - - - +

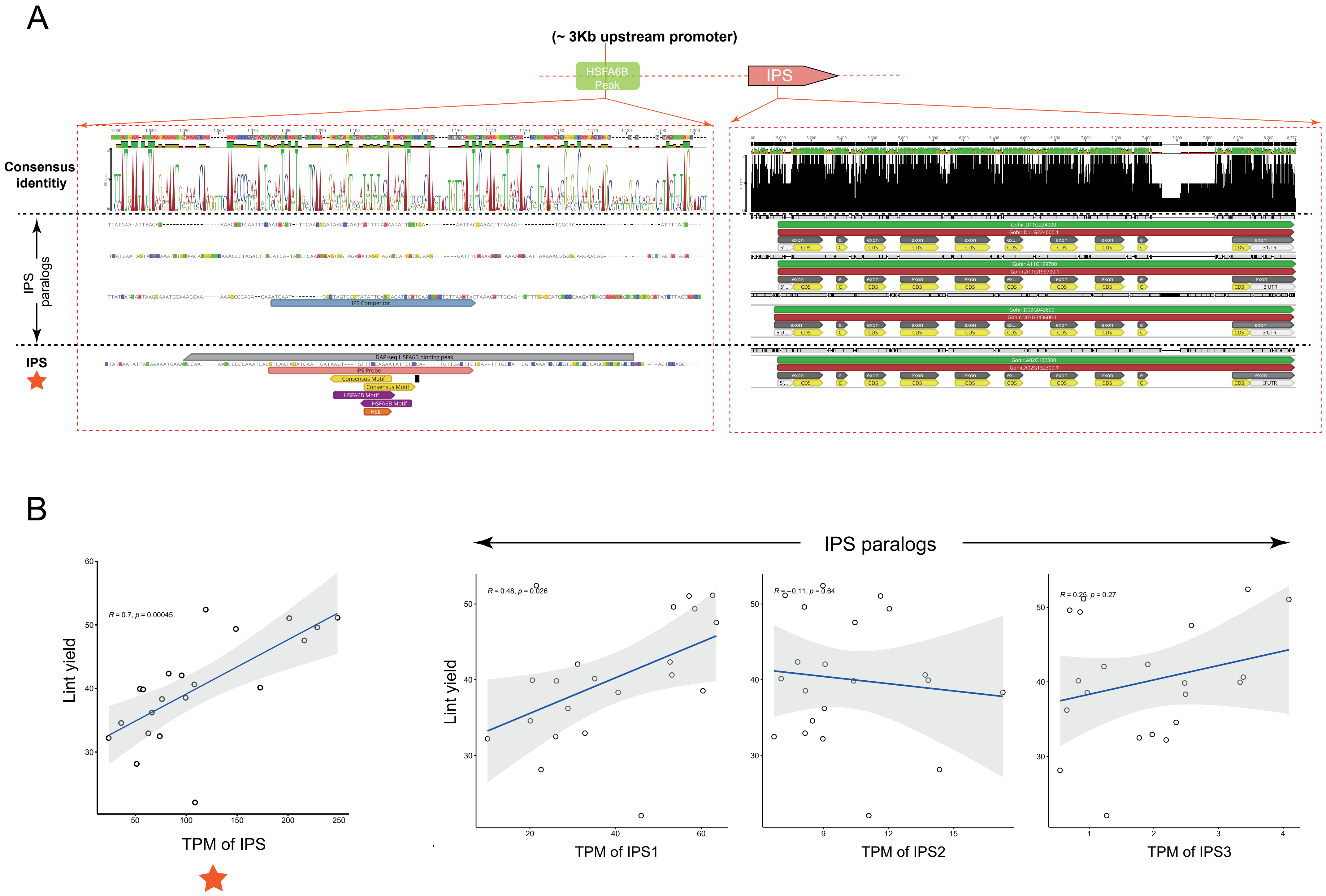
A
(~ 3Kb upstream promoter)
HSFA6B Peak
IPS
Consensus identitiy
IPS paralogs
IPS
DAP-seq HSFA6B binding peak
IPS Probe
IPS Competitor
Consensus Motif
HSFA6B Motif
HSE
Gohir.D11G224000
Gohir.A11G199700
Gohir.D03G043600
Gohir.A02G132300
B
IPS paralogs
R = 0.7, p = 0.00045
Lint yield
TPM of IPS
R = 0.48, p = 0.026
TPM of IPS1
R = -0.11, p = 0.64
TPM of IPS2
R = 0.25, p = 0.27
TPM of IPS3

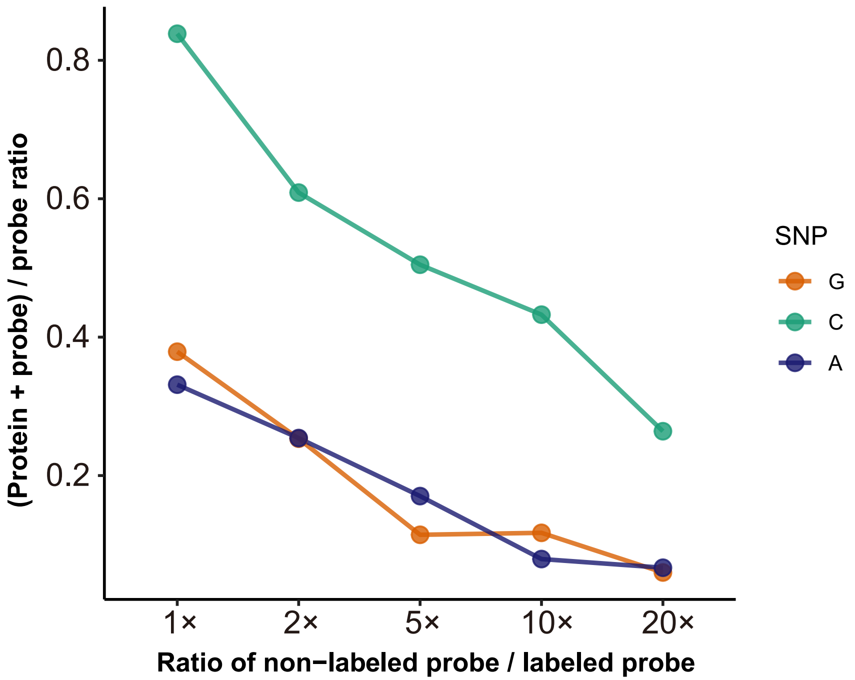

(Protein + probe) / probe ratio
0.8
0.6
0.4
0.2
1×
2×
5×
10×
20×
Ratio of non−labeled probe / labeled probe
SNP
G
C
A
