## Supplemental information for "Regulation of a single Inositol 1-Phosphate Synthase homeolog by HSFA6B contributes to fiber yield maintenance under drought conditions in upland cotton"

**Supplemental Information: EMSA Oligos used**

>ABP-Competitor-1

GGAAAATTCAAGCTCGATTCATGTTTAAGACGTTAAGATGAATAATGGAT

>ABP-Competitor-2

TTTTGGATTATTCTTGAAGCTTTCTGGAAAATTCAAGCTTGGATTCATGT

>ABP-Competitor-DREB2A

TCTCATGCAGCCAATGTCGTTGCTCCGCCACCGACTTAAATAATGGCACT

>ABP-Probe-1

GGAAAATTCAAGCTTGGATTCATGTTTAAGACGTTAAGATGAATAATGGA

>ABP-Probe-2

TTTTGGGGTATTCTTGAAGCTTTCTGGAAAATTCAAGCTCGATTCATGT

>ABP-Probe-DREB2A

TCTCATGCAGCCAATGTTGTTGCTCCGCCACCGACTTAAATAATGGCATT

>IPS-Competitor

ATCAATGGCTAGTGCGTATATTCATGACATCCATCAACTACTGTTAAATA

>IPS-probe

GTCAATAGATCAAGATAAGTTGTTTCCAGAATATTCCCCATGTTGACTTC

**Supplemental Information: EMSA Target sequences**

>Gohir.A12G006700-ABP-PromoterOnly

TCAGGGGTGAAATCAAGGGGACTAGTAGGGGCCTCTGTCTCTAAAAATAAAAAAATTTCCATTTAAATACTTTAACTCTCTAAAAATGATAAAAATTTGATTTAATCCTTTAAAATTTATAAAAATACAAATTATTAAAAAAATAAAATTACATTTTTACTATCGTAAACATTACGACTTAATTTCGTAAACATAATCATCATCACCACCTAGCATCTAACCCGAACCAAAATCTCCCATTTTCTTACAACCCACAACCTAAGTCACTATCATAGTCTTTTATCATTCTACCACTAAACCCAATCCCAACAATAAAATCTTCCTGTCAATAAAAAAAACCTGCAAGCAATAAGTAAACTGCAACAAAAATTCAAGCAACTCATTTAACTGTATCTTTAAAATCGAAAGCAACAGAGGAGAGAAAAAGAAGGCTCGAACCCAACCACATCTAAGGAGTTTTGGTTGTCCTTTTGCAAATCCGAATGGTGGAGCTGCAAAGGAGATTGCGAGGAAGCACTAGTTTTGAACGAAGGACAACTAGAGTGCCAACGTTGGAGATCGAATGAGAAGGAGCTGAGAGTTTGAGAATCATTTTGGATTATTCTTGAAGCTTTCTGGAAAATTCAAGCTTGGATTCATGTTTAAGACGTTAAGATGAATAATGGATCAATGCCATTATTTAAGTCGGTGGCGGAGCAACAACATTGGCTGCATGAGAGGAAGGAAATGTGTTTTTATGGTTTATTATGGAGACGAAGAAAGAAGAAAATTTTAAATATTTTATGTAAATAAAAAAAAAAAGTAAAGAGAAAGAAGGGTTTGGAGACCCATTTGCCGTAGAAAAGGAGAAAAAATTAATTAGGTGGAAATTAATAAATAACGTGGAATACAAATCAATACCTAAATTATATATATTTATAATTGGCTTGCACGCTCGTGATAAAGCAAATTACTTTCAACTCATACAATATAAGTATGACTATTCTTGTGATTCGAACTTACAACATGAGTTTTTAAGATAAGGGAGAGGAAACAATGGTGGAAGGAGGTCCAATGCAGGAGGGGGAGAGGAAGAGGAAGAGGGACAGGGACGAGGTGGGGGATTTTTATTTTACTTAATATTAAAATAATTATATTTATTGAAAATCTTTATGTTTTTTTTATAAATTTCTTTCAATTTTTTAGAATTAAGATTTAATTGATATCATTTGTAAATTTTGAGAGTTAATTTTTTTAAAATTATGACTAAATTGATATAATATGTAAAAGTTAAGGGCTAAAATTGTTATTATACCAAACTTTTTAATTGTATGTTAATTTGCTATTTGTGATTTTAATGTGAGTTGAGTACTTAAATTACAAACATTAATTTGGATGCTTAAATTTTATTTTTTTAATAATTAAGTGACTATATATATAATTTACCCTTAAGATAAACACCTAATACCACTAGACAAAAAGTCAAGGTTGAGTTTATAAAAATATATTAAGAAAATATTAAACTAAAACCATATAACTAAGGTTAACAAAATTGAAACAAAGATTTTCAAAATTGTAAGGTTATTTCTTTATTTTTTTTAAATTTAAAATATAAATCAAATTATTTAATATTGTTAAATTTTTTAAATTTTTTATTAAATTTATTGGTGTGAAATTTAAAATTCTAAAAAAATATTTATTTGATCTTCATGCAACAAAAAAAGTTTCGAACCTGAATATAATATCAACAATTGAACTTGAATTCTTCAATCTGAAAAGTCAAAAGACTAAATTATTCAAAATAAAAGTTGAGAGACTAAATTCTAAATTTACAAATAATACAGTGCTCAAAAATAAAATTTAACCATTTTGTTTTCTTTGTAGAACTTTCTGACGTGGCAGTTATGATGACATGGCAAAAACCCAAATGGGCCTATTCGGACACTCTTTGGGCTTTAGGCCCATTTTTTTAATTTAAAAAAGGAGTTATAAAAGACCAATTTCAGCCTATTCCGGAAAGTA

>Gohir.A12G006700-ABP-Gene-included

TCAGGGGTGAAATCAAGGGGACTAGTAGGGGCCTCTGTCTCTAAAAATAAAAAAATTTCCATTTAAATACTTTAACTCTCTAAAAATGATAAAAATTTGATTTAATCCTTTAAAATTTATAAAAATACAAATTATTAAAAAAATAAAATTACATTTTTACTATCGTAAACATTACGACTTAATTTCGTAAACATAATCATCATCACCACCTAGCATCTAACCCGAACCAAAATCTCCCATTTTCTTACAACCCACAACCTAAGTCACTATCATAGTCTTTTATCATTCTACCACTAAACCCAATCCCAACAATAAAATCTTCCTGTCAATAAAAAAAACCTGCAAGCAATAAGTAAACTGCAACAAAAATTCAAGCAACTCATTTAACTGTATCTTTAAAATCGAAAGCAACAGAGGAGAGAAAAAGAAGGCTCGAACCCAACCACATCTAAGGAGTTTTGGTTGTCCTTTTGCAAATCCGAATGGTGGAGCTGCAAAGGAGATTGCGAGGAAGCACTAGTTTTGAACGAAGGACAACTAGAGTGCCAACGTTGGAGATCGAATGAGAAGGAGCTGAGAGTTTGAGAATCATTTTGGATTATTCTTGAAGCTTTCTGGAAAATTCAAGCTTGGATTCATGTTTAAGACGTTAAGATGAATAATGGATCAATGCCATTATTTAAGTCGGTGGCGGAGCAACAACATTGGCTGCATGAGAGGAAGGAAATGTGTTTTTATGGTTTATTATGGAGACGAAGAAAGAAGAAAATTTTAAATATTTTATGTAAATAAAAAAAAAAAGTAAAGAGAAAGAAGGGTTTGGAGACCCATTTGCCGTAGAAAAGGAGAAAAAATTAATTAGGTGGAAATTAATAAATAACGTGGAATACAAATCAATACCTAAATTATATATATTTATAATTGGCTTGCACGCTCGTGATAAAGCAAATTACTTTCAACTCATACAATATAAGTATGACTATTCTTGTGATTCGAACTTACAACATGAGTTTTTAAGATAAGGGAGAGGAAACAATGGTGGAAGGAGGTCCAATGCAGGAGGGGGAGAGGAAGAGGAAGAGGGACAGGGACGAGGTGGGGGATTTTTATTTTACTTAATATTAAAATAATTATATTTATTGAAAATCTTTATGTTTTTTTTATAAATTTCTTTCAATTTTTTAGAATTAAGATTTAATTGATATCATTTGTAAATTTTGAGAGTTAATTTTTTTAAAATTATGACTAAATTGATATAATATGTAAAAGTTAAGGGCTAAAATTGTTATTATACCAAACTTTTTAATTGTATGTTAATTTGCTATTTGTGATTTTAATGTGAGTTGAGTACTTAAATTACAAACATTAATTTGGATGCTTAAATTTTATTTTTTTAATAATTAAGTGACTATATATATAATTTACCCTTAAGATAAACACCTAATACCACTAGACAAAAAGTCAAGGTTGAGTTTATAAAAATATATTAAGAAAATATTAAACTAAAACCATATAACTAAGGTTAACAAAATTGAAACAAAGATTTTCAAAATTGTAAGGTTATTTCTTTATTTTTTTTAAATTTAAAATATAAATCAAATTATTTAATATTGTTAAATTTTTTAAATTTTTTATTAAATTTATTGGTGTGAAATTTAAAATTCTAAAAAAATATTTATTTGATCTTCATGCAACAAAAAAAGTTTCGAACCTGAATATAATATCAACAATTGAACTTGAATTCTTCAATCTGAAAAGTCAAAAGACTAAATTATTCAAAATAAAAGTTGAGAGACTAAATTCTAAATTTACAAATAATACAGTGCTCAAAAATAAAATTTAACCATTTTGTTTTCTTTGTAGAACTTTCTGACGTGGCAGTTATGATGACATGGCAAAAACCCAAATGGGCCTATTCGGACACTCTTTGGGCTTTAGGCCCATTTTTTTAATTTAAAAAAGGAGTTATAAAAGACCAATTTCAGCCTATTCCGGAAAGTAACACTGAAAGATGACTGGACCTTGCTTCATTTTCTTCTTCCTTCTCTTAAACTTGCTTCCATTCTTTCGAACTCTCGAAGCTTCTCACTGCTCCATCAAAGGTAACCCTCTTTTCATTCCTCTTCCTTTCTAAGTTTGCTTCAGTCTTTCACCAATTTTCCCTCGTTTTCGAATCGGCTCGTTTTTGTTTCCACCATAGTTGCTTTATATTGACCTGGAGAAAAAAACATTGGAGATTGCAGCAGTTCCTTGATCATTGATTTATGCGGAGTTTACTTTATTTATTTGTTGAGTCAATACGGGTCAAGTTCATTAAGAAGATTACTAGGATTTCTTCAGTTTTTTAATGGCTGATTTTGTGTGAAATAATATATATTTTGGTGTTTTTTGATACAGGGTTACCTCTGGTGAGGAACATTGCTGATCTTCCACAGGATAATTATGGAAGAGGAGGTTTATCCCATATAACTGTTGCTGGTTCTCTCTTGCATGGGTTGAAAGAAGTATGCAGTTTATTTAGAGATATAATACTATTTCCTTTTTGTAAATTGGCCTACTTTTCTGAATTTTCACAAATAATTGTATTTTTATGTAAATTTGAATAGAGTTGAATATTCTCCATAGAATAACTTTTTTTGGGTTTTGCTTAGGTTGAGGTTTGGCTTCAAACATTTGCACCAGGATCGCGCACGCCGATCCATAGGCACTCTTGTGAAGAAGTTTTTGTTGTTCTCAAGGGCAGTGGCACTCTATATCTCGCCTCGAGTTCTAATAAGTACCCTGGAAAACCGGAGGAGCACTTTATATTTTCGAATAGCACGCTTCATATCCCTGTCAATGATGTTCACCAGGTAAAATATTTTCACTATTATGAATGTGATTGATGTGAAGAACCATAGGAATCTCGAAACAGAGACAGTTTTGTCGTAGGGTTCGCATGTAGTAAGATGATGTGGTTATTTATCACCTTTTGAAAAATAGTAGGAATCATGAAGCAACAAAATCCTCAGAGGATTAAGAAACTTTGATCATTTCCCTAGCCAATGCTATGCATCAATAATCGACATGATCGTATTAATCTTTACTTTATGAGGTTATGGAAAATGCATTGCCCCTTTCTTCACTTCAATGCTATCAAAGATGCTAATATCCTTCTATTTTTGTTTAAGTTTAACCTCCTTCTTAACTTTGGAAAACGTTCTTTCTGTTGGTGCAATCTAATCTTGCTTGGTAGTTGCCACTGATTTGGAGGATTGTTAATGTTTTCGTTTTAATTAGGTATCTCCACTAGATGCAAGTTTGGAGTGTGAATGTGCATTTCAGTTTAAGCACTAACATATGCTATAGTCTTTAGGTTAAAGATTTGATGACTGTAAGAATTACTATGACATTAGGTGAAAGGGAGGGAAGGACCCGATGAACAATGATGTCACTATGATGAATTTCACAGACGTTCCTTGTTTTTCAATTATTCTTGATTGCCATTGCAGGTCTGGAATACAAATGAACATGAGGATTTGCAAATGCTTGTGATAATATCTCGGCCGCCTATCAAAGTGTATGTAATTTTTCTTCTGTCTCGTTTTGCATCAACATTTTCACCATTAGTTGAGGGATGTCCTTTTATATTTCCTTTGTCCAGTTGAAAACCATGGTTGCTATCTTTGGCTTTCACCCTTCCAAATGTTTGCTGGTTGACACATTTTAGTTCTTGCTAGTTTCGAATGACATCTTCCTAACATTTAAAATAAACGAACTGAAAGCACCCATTCTCAAAGCACTTTGCCTTGGTTTTTATAGTCCTTTAAGTTGCGATAGTAGAGTACATCAAGATGGAATTTACTTACCTTGAAACTTTAACTACCCTTTACTCAATTAAGAGAAAGAAAAGTGAGAGCAGTTGTTTAGAAGCTTAAAGCAAAGGCTGTCCATGTATCTTTTAAAACATGACCCACATTTTCTTTCTCTAAATATGTTCCATTGCCATTGTTACTTGTTTAAGAACTCAAGAATGAACGACTGTTTTAGCATGCATGCATCATGATTGCAATCCCACCTGTTGATGTTTCCTGCATGATTGAAGATGTTGCTCAAGTTACTATATGCTAAATAGAATTTAAAGAACATTATTGGTTGCGATTGTTCATTCTTTTAACCCGACAACTGTATATTTTCAACAAATTTTCTTGTATGAAACTGAAAATGTATCCCTTGTTTTGCTAGTTCCTTGTGATACTTTTCCTTGATTCTCTCATTCATGAATCATGAGGATGATTTCTTTTTCTATTGAAATTTGCATTTAGGTTCATATATGAAGATTGGTTGATGCCTCACACTGCAGCTAAGTTGAAGTTTCCCTACTATTGGGATGAGCAGTGCTTTCAAGTACCTCAGAAAGATGAGCTTTAATTTTTGAAGACACGCCCCTTCACATGCTACTATATGAGCACTGTAATGGGGCCATTCCCATTTTACTGCTCAGATTACTTTACAAATTACATAAAGATTACAACATCTTAGCTTAGTTTGTATATTTTCCCCCTCATTTGAAGTCTGAATCCATTTTCTATTTTCATTTCATTTGACACTTCCAAAATCAATTGATGGTTTCCCTTGAAACTTCTCATCCCTCTGTTCCCAATGAATAGTTAAAATATGGCACTCTTCACAAATTAGGTGAAATTAAAATATAAGGATTAAATTTAAAAATTAATTTATAAAGCACACAAGTGGACAACAGTGTCTTGAAACTGAACCGGTCTGGTTCACCGTCAAAAACCA

>Gohir.D12G006100ABP-ABP-paralogs-PromoterOnly

TTTGTAGTGGTTTCATATATAATCTATAAATGGCACCCGTTTAAATTTTCTGATTTTATTGCCATTAATGGCTTTTTTTTACCGAGAAAAATAATGAATTTTTGGAAATATTTCATCATCACCACCTAGCATCTAACCCGAACCAAAATCTCCCATTTTCTTACAATCCACAACCCAAGGCACTATCATAGTCTTTATCATTCTACCACTAAACCCAAAACCTGCAAGCAATAAATAAACTGCAACAAAAATTCAAGCAACTCAGTTACCTGTATCTTTAAAATCGAAAGCAACAGAGGAGAGAAAAAGAAGGCTCGAACCCAACCGCATCTAAGGAGTTTTGGTTGTCCTTTTGCAAATCCGAACGGTGGAGCTGCAAAGGAGATTGCGAGGAAGCACTAGTTTTGAACGAAGGACGATTAGATTCCCTACGTTGGAGATCGAACAAGAAGGAGCTGAGAGTTTGAGAATCATTTTGGGGTATTCTTGAAGCTTTCTGGAAAATTCAAGCTCGATTCATGTTTAAGACGTTAAGATGAATAATGGATCAGTGCCATTATTTAAGTCGGTGGCGGAGCAACGACATTGGCTGCATGAGAGGGAGGAAATGTGTTTTTATGGTTTACTATGGAGACGAAGAAAGAATAAAATTTTAAATATATTTTTATGTAAATAAAAAAAAATGGTAGAGAGAAAGAAGGGTTTGGAGACCCATTTGCCGTAGAATAGGAGAAAAAATTAATTAGGTGGGAATTAATAAATGACGTGGAATGCAAATCAATACCTCATTAAGGAGAAAGCGGACGAGTGATGATTTGCACAATTTAATTAAAAAATTTTAAGTGATCAAAATAGGAATATTTCATTTTTAGGGTGACTAATGGGATATTTTACCCAAATTATATATATTCATAATTGACTTGCACACACATGACAGAAGTAAATCACTATCAACTCACACACACAAGAAATATGACTATCCTTGTGATTCGAACTTACAACATGAGTTTTTAAGACAAGGGAGAGGAAACAATGGTGGAGGGAGGTACAATGGAGGAGGGGGAGAGGAAGAGGAAGAGGGATGAGGTTGGGGATTTTTATTTTATTTAATATTAAAATAATTATATCTATTGAAAACATTTATGTTTTTTTTATAAATTTCTTTCAATTTTTTAGAATTAGGATCTAATTGAGATCAATTGTAAATTTTGAAGGTTAATTTTTTAAAAATTATGACTAAATTGATATAATATGTAAAAGTTGAGGGCTAAAATTGTTATTATACCAAAATTTTTAACTGTCATGTTAACCTGCTATTTGTGATTTTAATGGAGTGACCAAATGAGTACTTAAATTACAATCTTTTAATTTGGGTACTTAAATTTTATTTTTTAATAATTAGGTGACTATATATACAATTTACCCTTAAGATAAACACCTAATACCGCTAGACAAAAAGTCGAGGTTGAGTTTATAAAAATATAAATATTAAACTTGTTTAAATAGTGATTGGCATAAAATGTTACTAAAACCATATAACGAAGGTTAACAAAATTGAAACAAAGATTTTTAAAATTGTAAGGTTGAATTTAATTTCTTTATTTTTTAAAATTTTAAAATATAAATAAAATTATTTAATATTGTTAATATTTTAAAATTTTTATTAAATTTATTGGTGTGAAATTTTAAAAATTTTTAAAAAATATTTATTTGTTAGTCATGCAACAAAAAAAGTTTCGAACCTGAATATATATGAAAAAAAGATTAAATTATTCAAAATAAAAGTTGAGAGACTAAATTCTAAATCTACAAAGAATACAGTGCTCAAAATAAAATTTAACCATTTTGTTTTCTTTGGAGAACTTTCTGACGTGGCAGTTATGATGACATGGCAAAAACCGAATGGGCCTATTCGGACACTCTTTGGGCTTTAGGCCCATATTTTTTAATTGAAAAAAGGAGAAATAAAAGACCAATTTCAGCCTATTCCGGAAAGTAA

>Gohir.D12G006100ABP-ABP-paralogs-Gene-included

ATTTGTAGTGGTTTCATATATAATCTATAAATGGCACCCGTTTAAATTTTCTGATTTTATTGCCATTAATGGCTTTTTTTTACCGAGAAAAATAATGAATTTTTGGAAATATTTCATCATCACCACCTAGCATCTAACCCGAACCAAAATCTCCCATTTTCTTACAATCCACAACCCAAGGCACTATCATAGTCTTTATCATTCTACCACTAAACCCAAAACCTGCAAGCAATAAATAAACTGCAACAAAAATTCAAGCAACTCAGTTACCTGTATCTTTAAAATCGAAAGCAACAGAGGAGAGAAAAAGAAGGCTCGAACCCAACCGCATCTAAGGAGTTTTGGTTGTCCTTTTGCAAATCCGAACGGTGGAGCTGCAAAGGAGATTGCGAGGAAGCACTAGTTTTGAACGAAGGACGATTAGATTCCCTACGTTGGAGATCGAACAAGAAGGAGCTGAGAGTTTGAGAATCATTTTGGGGTATTCTTGAAGCTTTCTGGAAAATTCAAGCTCGATTCATGTTTAAGACGTTAAGATGAATAATGGATCAGTGCCATTATTTAAGTCGGTGGCGGAGCAACGACATTGGCTGCATGAGAGGGAGGAAATGTGTTTTTATGGTTTACTATGGAGACGAAGAAAGAATAAAATTTTAAATATATTTTTATGTAAATAAAAAAAAATGGTAGAGAGAAAGAAGGGTTTGGAGACCCATTTGCCGTAGAATAGGAGAAAAAATTAATTAGGTGGGAATTAATAAATGACGTGGAATGCAAATCAATACCTCATTAAGGAGAAAGCGGACGAGTGATGATTTGCACAATTTAATTAAAAAATTTTAAGTGATCAAAATAGGAATATTTCATTTTTAGGGTGACTAATGGGATATTTTACCCAAATTATATATATTCATAATTGACTTGCACACACATGACAGAAGTAAATCACTATCAACTCACACACACAAGAAATATGACTATCCTTGTGATTCGAACTTACAACATGAGTTTTTAAGACAAGGGAGAGGAAACAATGGTGGAGGGAGGTACAATGGAGGAGGGGGAGAGGAAGAGGAAGAGGGATGAGGTTGGGGATTTTTATTTTATTTAATATTAAAATAATTATATCTATTGAAAACATTTATGTTTTTTTTATAAATTTCTTTCAATTTTTTAGAATTAGGATCTAATTGAGATCAATTGTAAATTTTGAAGGTTAATTTTTTAAAAATTATGACTAAATTGATATAATATGTAAAAGTTGAGGGCTAAAATTGTTATTATACCAAAATTTTTAACTGTCATGTTAACCTGCTATTTGTGATTTTAATGGAGTGACCAAATGAGTACTTAAATTACAATCTTTTAATTTGGGTACTTAAATTTTATTTTTTAATAATTAGGTGACTATATATACAATTTACCCTTAAGATAAACACCTAATACCGCTAGACAAAAAGTCGAGGTTGAGTTTATAAAAATATAAATATTAAACTTGTTTAAATAGTGATTGGCATAAAATGTTACTAAAACCATATAACGAAGGTTAACAAAATTGAAACAAAGATTTTTAAAATTGTAAGGTTGAATTTAATTTCTTTATTTTTTAAAATTTTAAAATATAAATAAAATTATTTAATATTGTTAATATTTTAAAATTTTTATTAAATTTATTGGTGTGAAATTTTAAAAATTTTTAAAAAATATTTATTTGTTAGTCATGCAACAAAAAAAGTTTCGAACCTGAATATATATGAAAAAAAGATTAAATTATTCAAAATAAAAGTTGAGAGACTAAATTCTAAATCTACAAAGAATACAGTGCTCAAAATAAAATTTAACCATTTTGTTTTCTTTGGAGAACTTTCTGACGTGGCAGTTATGATGACATGGCAAAAACCGAATGGGCCTATTCGGACACTCTTTGGGCTTTAGGCCCATATTTTTTAATTGAAAAAAGGAGAAATAAAAGACCAATTTCAGCCTATTCCGGAAAGTAACAGAGAAAGATGACTGGACCTTGCTTCATTTCCTTCTTCCTTCTCTTAAACTTGCTTCCATTCTTTCAAACTCTCGAAGCTTCTCACTGCTCCATCAAAGGTAACCCTCTTTTCATTCCTCTTCCTTTCTGGGTTTGCTTCAGTCTTTCACCAATTTTCCCTCGTTTTCGAATCGGCTCGTTTTAGTTTCCACCATAGTTGCTTTATATTGACCTGGAGAAAAAAACATTGGAGGTTGCAGCAGTTCCTTGATCATTGATTCATGTTGAGTTTAATTTATTTATTTGTTCAGTCAATACGGGTCAAGTTCATTAAGAAGATTACTAGGATTTTTTCAGTTTTTTAATGGCTGATTTTGTGTGAAATAATATATATTTTGGTGTTTTTTGATACAGGGTTACCTCTGGTGAGGAACATTGCTGATCTTCCACAGGATAATTATGGAAGAGGAGGTTTATCCCACATAACTGTTGCCGGTTCTCTCTTGCATGGTTTGAAAGAAGTATGCAGTTTATTTAGATATATAATACTATTTCCTTTTGTAAATTGGCCTACTTTTCCGAATTTTCACAAATAATTGTATTTTTATGTAAATTTGAATAGAAGATGTTGAATATTCTCCATAGAATAACTGTTTTTGGGTTTTGCTTAGGTTGAGGTTTGGCTTCAAACATTTGCACCAGGATCGCGCACGCCGATCCATAGGCACTCTTGTGAAGAAGTTTTTGTTGTTCTTAAGGGCAGTGGCACTCTATATCTCGCCTCGAGTTCTAATAAGTACCCTGGAAAACCGGAGGAGCACTTTATAATTTTCGAATAACACGTTTCATATCCCTGTCAATGATGTTCACCAGGTAAAATATTTTCACTACTATGAATGTGATTGATGTGAAGAACCATAGGAATCTCGAAACAGAGACAGTTTTGTCTTAGGGTTCGCATGTAGTAAGATGATGTGGTTATTTATCACCTTTTGAAAAATAGTAGGAATCATGAAGCAAGAAAATCCTCAGAGGATTGAGAAACTTTGATCATTTCCCTAGCCAATGCTATGCATCAATAATCAATATGGTCGTATTAATCTTTACTTTATAAGGTTATGGAAAATGCATTGCCCCTTTCTTCACTTCAATGCTATCAAAGATGCTAATTGCTAATATCCTTCTATTTTTGTTTAAGTTTAACCTCCTTCTTAACTTTGGAAAACGTTCTCTTTGTTGGTGCAATCTAATCTTGCTTGGTAGTTGCCACTGATTTGGAGGATTGTTAATGTTTTCGGTTTAATTAGGTATCTCCACTAGATGCAAGTTTGGAGTGTGAATGTGCATTTCAGTTTAAGCACTAACATATGCTATAGTCTTTAGGTTAAAGATTTGATGACTGTAAGAATTACTATGATACTAGGTGAAAGGGAGGGAAGGACCCGATGAACAATGACGTCACTATGATGAATTTCACAGACGTTCCTTGTTTTTCAATTATTCATGATTGCCATTGCAGGTCTGGAATACAAATGAACATGAGGATTTGCAAATGCTTGTGATAATATCTCGGCCTCCTATCAAAGTGTATGTAATTTTTCTTCTGTCTCGTTTTGCATCAACATTTTCACCATTAGTTGATGGATGTCCTTTTGTATTTCCATTGTCCAGTTGAAAACCATGGTTGCTATCTTTGGCTTTCACCCTTCCAAATGTTTGCTGGTCGACACATTTTAGTTCTTGCTAGTTTCGAATGACATCTTCCTAACATTTAAAATAAACGAACCGAAAGCACCCATTCTCGAAGCACTTTGCCTTGGTTTTTATAGTCCTTTAAGTTGCTATATTAGAGTACATCAAGATGGAATTTACTTACCTTGAAACTTTAACTGTCCTTTACTCAATTAAGAGAAAGAAAAGTGGGAGCAGTTGTTTGGAAGCTTACAGCAAAGGCTGTCCATGTATCTTTTAAAACATGACCCACATTTTCTTTATCTAAACATGTTCCATTGCCATTGTTACTTGTTTAAGAACACAAGAATGAATGACTGTTTGAGCATGCACGCATCATGATTGCAATTCCACCTGTTGATGTTTCCTGCATGATTGAAGATGTTGCTCAAGTTACTATACGCTAAATAGAATTTAAAGAACATTATTGGGTGCGATTGTTCATTCTTTTAACCCGAAAACTTTATATTTTCATTTTCTTGTATGAAACTGAAAATGCGGCCCTTGTTTTACTAGTTCCTTGTGATACTTTCCCTTGATTCTCTCATTCATGAATGCTGAGGATGATTACTTTTCCTTTTTCAATTGAAATTTGCATTTAGGTTCATATATGAAGATTGGTTGATGCCTCACACTGCAGCCAAGTTTGAAGTTTCCCTACTATTGGGATGAGCAGTGCTTTCAAGTACCTCAGAAAGATGAGCTTTAATTTTTGAAGACACGCCCCTTCACATGCTACTATATGAACACTGTAATGGGGCCATTCCCATTTTACTGCTCAGATTACTTTACAAATTACATAAAGATTACAACATCTTAGCTTAGTGTGTATATTTTCCCACTCATTTTGAAGTCTGGATCCATTTTCTATTTTCATTTCATTTGACACTTTCATAATCAATTGATGGTTTCCCTTGAAACTTCTCATCCCTCTGTTCCCAATGAATAGTTAAAATATGGCACTCTTCACAAATTAGGTGAAATTAAAATATAAGGATTAAATTTAAAAATTAACATAGTTGAGGGATTCAAATCAAAATTAGACCTTTTATTTATAAGCAAACAAGTGGACAACAGTGTCTAGAAACTGAACCGGTCTGGTTCACC

>Gohir.A02G132300-IPS-Gene-included

TACGAGGCCCAACCCATCGTAACTTGTTTTTCTCTCATGTAACCGATCTTCCCAAGTCTATCTTCGAAAGTCGATTCTTAAGTAAATCAACCACGTCTATCGCCGTTAAAATACTTAGTTTTAAAACAGGAAAAGCTCAACTCGGAAGTTGTTTTATCAAGAATCATTTCTGATCTCGTCTTTCCTTAGCGGGCCGCCGCTAAGAGTGTTCATAGGAAGTCGAAATGCCACAGGCCGATCCTTCCTCCCAACTGTAAAATTAAACGATTCCAGACCAACAGATCCTTTTTGATTTGAAAAACGTTTAAAAATTTTAGTAAGGCCGATTGTCAAACTATCTAAGTAGAAAGACGAAAATGTATCCCGAAGGGATTTAAGCAATATTCAATTCTCACGTGAGGAATTGATTCGTACTAGCTCTTTGCTTGAGAAAAATTAGCCAAGCATGATATTGATTATCATGCTTAACATGTTTTGATAATGTAAAAACATAAAATAAAATAAAGTAAAGTAGAAATAAAACAAAGAAGAAAATTTTATTAAAAATGAAAACAAAGTAGCAATAAGAACAAAAGACAAGAGATACTTAGCCTTACCCAAATGTGGGTATTTATAGGCTTCAAATTCCCAAATCAACAAGGAAAACTAATCAAAAAAAGATAAGATAATATCTATAAAATTATGAAAATGAAATCCAAAATCCCCCAAATCACGTCAATAGATCAAGATAAGTTGTTTCCAGAATATTCCCCATGTTGACTTCTGATTTGCCACGTCAAATCCGCGTCGTGCCCAGCGTCGTGCCCACGACTCCAGCCGTGACAGAATAAGCGTCGTGTCCACGACTCCAACCGTGACAGAATAAAAAGAAAAAAAAACTGCAGATTACGTCTTTTTGTTGCCATTTCTAGTATTGGGTTCACTACACAAGAAATAAATCAAATATAGCCAATTAAGTAGTGAAAAATTCATAAGACAAGGGAATTAACAAGCAAAAAGTGTCAAATATAAGAGTACATCAGGTTGTTGATCATATTGCAAAAATAGCTTCAAGTGAATTGGGAAACACACAACTATTTTGGGAGCATCTAGATTCAATAAAGAATATACTGATAACTGAGTGCAATACTATTTCTCTATGTCGATAGTACTTTAAGAATGTAATCACTTTGTCTTGTTATTTTCTACCGAAACAAATTGAACTGACTAATATAACTTTGTGTTTTTCTTTTTTACCATTCTAAATATACAAACTTATAAGATTTCTAGTTTTACTAAAATAATTGGTTAAAAAATTACTTGATCAAATTTATTATAATTATAAATGAGAATTTTAATTCTCTTAAATGACCTTTTTAAAGGAATTTATATAATAATAATAATAAGTCAAATTTATATTGAATTTTTTTAAATTTAAATTATTTTGCTTCCATTAACCAAACAGTGAAAAGTGAAAAATTATATTACTTTTTAAATAGTAAATTTTAATTATTTATAAAATAATATTTTATATATAAAATTTTATAAATAATACATTTTTAATATAATATAATATATTAATTAATATATTAATTATATATTATCTGTAAAATGTTTTGGGAATATATTAACGGCTAAGAAAAAAATAAAAAAATAAAAAATTAAGAATTTAATTTACTATATTAATTTTTATAATTTAAAAATTACATGAAGTAAAACTACACTATCAATTGAATATGTTAAAAATATTTTAATTATTGTTTCAATTTATTATTATACTATTTAGATTTTAAATAAATTAATATATAAAATTACATACAATACATATGAGATTAGAGACTAATATATATTCTGAAAAATAAATGCTCTTTATTATTTATTATACTTCAAAATTGCCATTATAGTTTTTATTACATTTGAAATTTATCAAATGTATAACTAAAAAATTTTAATTTTTTCGAAACGCTTTTTTAATGAATTGAATTTGATAATAGAAAAAAGATATTTTAAAAAAAATCATTACATATTATATTTATTTTAAGTGCTTTAGAGGATTTTTATTTCTAATAATGTTTATCAAAATTATAACTAACACATTTAAATTTATCAAAAATTTGTTAAGAAATTAATTTAATAAATTTTTTTGACTCATCGAAAAGATAAATTTTAAACTTTTTTAAAAACATGTTAATTTATAAACAATTAAGCATAATTAAATAATAAGTTTTAAGAGATTATTATTTTTAACAATGATTAAAAAATAATAATATTTTGAAAAGTTAGCTTAAATATCTAACATAATTATTGTGAACAATAAAATATATTTTAATATGAAAAAAAAGTACATTATAAAAGTTTCATGAAAAACAATATATTATATTCTACCAAAGCACGAGATACACTTAGTGGCATTAAAAAATTGTTAACTAAAATACAAAAATAATTTAATATTTTTATAAATTAGCAATTTGAGAAAACTCTAATCTTGATCTCAGTCTGGTTTATTTTTAAGTGTCAAAGTCTCAGTCAAATCCTTTTATTTATTTACGACCCTGATTCTAGCTGACAAGTACGGGATCTCATCTTTTTAACCATGCCAGATTAATAGTTAAACTGTAAATTTTGGTTTTTTTATCCTAATAATAATAAAAATAACTTAAATAATATGCTCTTAAGATTATTTATCAAAATGATATCTTTATTTCAATATTTTACTGGTGTTACCTAATAAATTGGCGACACCATACTGAAACATGATGACTTAAAATATTCAAAATCTAATATATAAGTTGACAGTGCGGCGACACCAAACTTTAAATAAAGTAAATTGTTTTTTAAACTCTTTTTGTTTATTATTTCTTTTTATTTCTCTTTAACACAAAAAATAGAGAGACTTTTTATCAATTAGTCTATAAAGTTTTTTTTACTAAAAAGTATTTTTGTAAAAAAATCGATTTTAACTAGAAATAAATTTTATTTATTTTTAAAAAAATAATTTTTTTGTTCAAATAATTTTTTTTAAAAATTATTCCGACAAGAATAAAAGAATTTGAACAAAAATTATTTTTCAAAAAATAAATAAAATTTATTTTCAATAGGAATCAATTTTTTTACAAAAAGAAAATTTGATAGAAATTTTTTTTGAGTAATTGGTAAGAAGCCTCTCTATTTTTTGTATTAAAAGAAAATAAAAGAAAATAATAAATAGAGAAAGTTTATGAAGCAGTTTAACTTATTTAAAGTTTGATGCTCTCACACTATTAACGGACCTAACAATTCTTTTTAACAACCAATCAAAATGAGAAATTTACGTATTAAATCCCTGAATATTTTAAGTCAATATATTTTAATCTTGTGTCACTAATTTACCCCAACACTGACAAAATACTGTAGTAAAAGTACTATTTTGATAAATAATTTTATGGGTATATATTTGAATTATTAAATTTTTTTATATTATTGATGTAAGAAAAGCCGTATCTTTTATTATTATTACTTCCCACTCTGGCCTCACATGAAAACAAGCAAGGACGGACTCAGTTGATAGTTGAGATGCACAATACACCTCAACTAAAGCGCGTAAGAAACACCTATTCTTCACCGTCGGCCAAGTCTTCGGCATATACCATTAAAATTCCGGTCTTTTATCCACGTGGAAATACAGCGCGTGGATATGGGTCCTACTAAACGATCCAGTCATTCAATCAATATACCCCTTCGAGTACCTTCTCAATGGCAACACAATAAAAATAAAATAAAATAAAATTTCCTTCACAAGGTTAAGCCACGTGTAAAGTGTATTACACGTCAAAAATGACACCCCTCTTCACACGTACGAACGGGCAAAAAAACCACAAAACAGCATAGGCATGGCCTACCTTCTCTTGAATCGAAAAACCAAAGGGGATTCAATGACGAAATTTTAACGTAAGGCATCAACCATTGTGATGATGCCACGTGTATCCTCTCATTATATGTTTCCTCTATAAATAAGCCTCCTAGTCCCAAGCTTTGAAGCACAAAATTGTGGGGCAAAAGAGAAATAAAAACAGTTTCATTTCCATTCTAAGTTTGTGTTTTTTATTTGCCTTTTGTGGGTTTTTTTTAAAAAAAATTCAGCAAAAATGTTTATTGAGAGTTTTAAGGTAGAGAGCCCCAATGTGAAGTACACGGAGAATGAAATTCAGTCTGTTTACAACTATGAAACTACAGAGCTTGTTCATGAGAACAAAAATGGAACCTATCAATGGGTTGTTAAACCCAAAACTGTCAAATATGAATTCAAGACTGATATCCATGTCCCTAAATTGGGGTTAGAATCCAAATCTGCCCATCATTTGTTTTTTTGCCATTTCGATTTTGATTATTTTGATTTTATTTATTTATGTGTTTTAGGGTGATGCTTGTGGGATGGGGAGGAAACAATGGTTCAACCCTCACCGGAGGTGTTATAGCTAACAAAGAGTGAGTTTTGTACTTTCTAGATAAAAGATCTGATTAATTTTGATTTGTTTTCTTTTCTCCTTTTTTTTTTTTGGATTTTTGAGATATGGGTAATGATGGGTTTTCTCAGGGGTATCTCTTGGGCTACTAAGGACAAGGTACAACAGGCTAATTACTTTGGTTCATTGACTCAAGCATCAACGATCCGAGTTGGGTCCTACAATGGAGAAGAGATTTATGCTCCATTTAAGAGTCTTCTTCCTATGGTATACGAATTATAAACCCAGATTTAGTTTTTAATTTACTAGTTTGCTACTTAATGCTTAATGCTTGATAAATGAATGAATAAAGGTGAACCCAAATGATATTGTGTTTGGAGGATGGGACATTAGTGACATGAACCTAGCTGATGCAATGGCTAGGGCCAAGGTTTTCGACATTGATCTGCAAAAGCAACTGAGACCCTACATGGAATCCATGGTCCCACTCCCTGGAATCTACGATCCTGATTTCATTGCTGCTAACCAAGGTGAACGTGCCAATAATGTCATGAAGGGGACCAAGAAAGAACAAGTTCAGCAGGTCATCAAAGACATCAGGTATATAATCAACTCCAACCCTCCAAACTGTATTTGTTAAGCTTGTAGAACAGGAAAAGTAGCTGAATTTTTTAACATTTGATTGGGTAAATCAGGGAGTTCAAAGAGAAAAACAAGGTGGACAAGGTTGTTGTACTCTGGACTGCAAACACTGAGAGGTACAGCAATGTCATTGTGGGGCTAAATGACACCGTGGAAAGCCTTATGGCTTCTTTGGAGAAGAATGAATCAGAGATTTCTCCTTCCACTTTGTATGCTATTGCTTGTGTTCTTGAAAATGTTCCTTTCATCAATGGCAGCCCACAAAACACCTTTGTTCCAGGTTCTTAATTTAGCTCAGCTGTCTTCCTAATTTTATAAATTGTAATGACGAGTAAATGTAATAACCTCTACAAGTGACTCTTATTGTATTCAGGGTTGATTGATTTGGCTATTCAAAGGAACTGTCTGATTGGAGGAGATGACTTCAAGAGTGGCCAGACCAAGATGAAATCTGTCCTCGTGGATTTCCTTGTTGGGGCTGGTATCAAGGTATAAATCTTTTGACTTTATGGCACTATCGCCGTAATTGTTATAATAAGCAATTAGTACCAATATAATTGAAGTAGAAAGTTTAAGTGTTTGGTAGCTTATGTGTCAATAAAATGTAAATAATATGACATTGATAGGTGGTTGTACTAATGGTGATGATAAATTGACAGCCAACATCGATAGTGAGTTACAACCATCTGGGAAATAATGATGGCATGAATCTGTCAGCACCCCAAACCTTCCGTTCCAAGGAGATCTCAAAGAGCAATGTTGTTGATGACATGGTTTCAAGCAATGGAATCCTCTATGAGCCTGGTGAACATCCTGATCATGTTGTGGTCATCAAGGTAAATATTTTTTTACCCATCTGTGTCTGTTTCGGCACTTGTGAGTGGTGACCTTAAAGCTTTTTTGCTTGATTGGTCTTAAAATGTGTACTTTGTCTTCCTTGTATTATAGTATGTGCCGTATGTGGGAGACAGCAAGAGAGCCATGGATGAGTACACATCAGAGATATTCATGGGAGGCAAGAACACCATTGTGTTGCACAACACATGTGAGGATTCCCTGTTGGCTGCTCCCATTATCCTAGACTTGGTTCTCCTTGCTGAACTTAGCACCAGGATCCAGTTCAAGGCTGATGGAGAGGTATGACCCTTAGAACTTTGTGGACTCTTCTGGGTTGACCCTTAAAACATATTTATGCAAAAACATTGATGTTGAATTTTGGTTTTGCAGGGCAAGTTCCACACTTTCCATCCTGTGGCTACAATCCTCAGTTACCTCACCAAAGCCCCTCTTGTGAGTCCAAAGCACCCCTTTGTCTTTTCCTGTAACTCTGGCCTACTAAAAATATAATAATAATTAATAATAAAACGGGGGTGGGGAGGGTTGACACCTTCACCAATAAAACATTGATATTATTGTCTTGAATTAAATGTTAACTGTTTATGGTTTAAGTGGGAAACAAGGAATCGCAAATTCACTCTATGGTGCCTCTTTGTGATATATTTCATGTGACCATTAATGATTTTATGTAAAGCTGCTTCTGTCTATTTCACGTGAAGCAACTGGTGAGGGGGGACACAATATTGAATTAAAGTTTGTGAACTTTTATGTCAAATTTCAGTTCAAGAATGTGAATAGAATCATGAGAATCACTTGAATCAGAAGTCAAGTGATTGTGAAGAAATTTACTGAGAACTGACATTGTTTTGGCAAAAAACAGGTTCCACCAGGCACACCGGTGGTGAACGCACTGTCCAAGCAGCGTGCAATGCTGGAGAACATACTAAGGGCCAGCATTGGCTTGGCTCCTGAAAACAACATGATTTTGGAATACAAGTGAAAAAGTTGAGAGAAAAGTGGAACAAAGCTTTGGTTCTTAGAACTTTAGGACCATTTTCCCTTTCTCTGTCGTTTTCATAAAAGCTGTGGTTACTGTTGTCACAGATTGTAACGGTTGTATGATGGTTTATGCTATGCTTCTTTTTTTCTCATTTATTTTCAAGTGGCGCTCATGTTTGTAGCTCACTCTTGAGCCAAGCACCTCTTTTGTGGAAATAGATATGATAGTATGAATTTTCGTAACTGGACAATCT

>Gohir.A02G132300-IPS-PromoterOnly

TACGAGGCCCAACCCATCGTAACTTGTTTTTCTCTCATGTAACCGATCTTCCCAAGTCTATCTTCGAAAGTCGATTCTTAAGTAAATCAACCACGTCTATCGCCGTTAAAATACTTAGTTTTAAAACAGGAAAAGCTCAACTCGGAAGTTGTTTTATCAAGAATCATTTCTGATCTCGTCTTTCCTTAGCGGGCCGCCGCTAAGAGTGTTCATAGGAAGTCGAAATGCCACAGGCCGATCCTTCCTCCCAACTGTAAAATTAAACGATTCCAGACCAACAGATCCTTTTTGATTTGAAAAACGTTTAAAAATTTTAGTAAGGCCGATTGTCAAACTATCTAAGTAGAAAGACGAAAATGTATCCCGAAGGGATTTAAGCAATATTCAATTCTCACGTGAGGAATTGATTCGTACTAGCTCTTTGCTTGAGAAAAATTAGCCAAGCATGATATTGATTATCATGCTTAACATGTTTTGATAATGTAAAAACATAAAATAAAATAAAGTAAAGTAGAAATAAAACAAAGAAGAAAATTTTATTAAAAATGAAAACAAAGTAGCAATAAGAACAAAAGACAAGAGATACTTAGCCTTACCCAAATGTGGGTATTTATAGGCTTCAAATTCCCAAATCAACAAGGAAAACTAATCAAAAAAAGATAAGATAATATCTATAAAATTATGAAAATGAAATCCAAAATCCCCCAAATCACGTCAATAGATCAAGATAAGTTGTTTCCAGAATATTCCCCATGTTGACTTCTGATTTGCCACGTCAAATCCGCGTCGTGCCCAGCGTCGTGCCCACGACTCCAGCCGTGACAGAATAAGCGTCGTGTCCACGACTCCAACCGTGACAGAATAAAAAGAAAAAAAAACTGCAGATTACGTCTTTTTGTTGCCATTTCTAGTATTGGGTTCACTACACAAGAAATAAATCAAATATAGCCAATTAAGTAGTGAAAAATTCATAAGACAAGGGAATTAACAAGCAAAAAGTGTCAAATATAAGAGTACATCAGGTTGTTGATCATATTGCAAAAATAGCTTCAAGTGAATTGGGAAACACACAACTATTTTGGGAGCATCTAGATTCAATAAAGAATATACTGATAACTGAGTGCAATACTATTTCTCTATGTCGATAGTACTTTAAGAATGTAATCACTTTGTCTTGTTATTTTCTACCGAAACAAATTGAACTGACTAATATAACTTTGTGTTTTTCTTTTTTACCATTCTAAATATACAAACTTATAAGATTTCTAGTTTTACTAAAATAATTGGTTAAAAAATTACTTGATCAAATTTATTATAATTATAAATGAGAATTTTAATTCTCTTAAATGACCTTTTTAAAGGAATTTATATAATAATAATAATAAGTCAAATTTATATTGAATTTTTTTAAATTTAAATTATTTTGCTTCCATTAACCAAACAGTGAAAAGTGAAAAATTATATTACTTTTTAAATAGTAAATTTTAATTATTTATAAAATAATATTTTATATATAAAATTTTATAAATAATACATTTTTAATATAATATAATATATTAATTAATATATTAATTATATATTATCTGTAAAATGTTTTGGGAATATATTAACGGCTAAGAAAAAAATAAAAAAATAAAAAATTAAGAATTTAATTTACTATATTAATTTTTATAATTTAAAAATTACATGAAGTAAAACTACACTATCAATTGAATATGTTAAAAATATTTTAATTATTGTTTCAATTTATTATTATACTATTTAGATTTTAAATAAATTAATATATAAAATTACATACAATACATATGAGATTAGAGACTAATATATATTCTGAAAAATAAATGCTCTTTATTATTTATTATACTTCAAAATTGCCATTATAGTTTTTATTACATTTGAAATTTATCAAATGTATAACTAAAAAATTTTAATTTTTTCGAAACGCTTTTTTAATGAATTGAATTTGATAATAGAAAAAAGATATTTTAAAAAAAATCATTACATATTATATTTATTTTAAGTGCTTTAGAGGATTTTTATTTCTAATAATGTTTATCAAAATTATAACTAACACATTTAAATTTATCAAAAATTTGTTAAGAAATTAATTTAATAAATTTTTTTGACTCATCGAAAAGATAAATTTTAAACTTTTTTAAAAACATGTTAATTTATAAACAATTAAGCATAATTAAATAATAAGTTTTAAGAGATTATTATTTTTAACAATGATTAAAAAATAATAATATTTTGAAAAGTTAGCTTAAATATCTAACATAATTATTGTGAACAATAAAATATATTTTAATATGAAAAAAAAGTACATTATAAAAGTTTCATGAAAAACAATATATTATATTCTACCAAAGCACGAGATACACTTAGTGGCATTAAAAAATTGTTAACTAAAATACAAAAATAATTTAATATTTTTATAAATTAGCAATTTGAGAAAACTCTAATCTTGATCTCAGTCTGGTTTATTTTTAAGTGTCAAAGTCTCAGTCAAATCCTTTTATTTATTTACGACCCTGATTCTAGCTGACAAGTACGGGATCTCATCTTTTTAACCATGCCAGATTAATAGTTAAACTGTAAATTTTGGTTTTTTTATCCTAATAATAATAAAAATAACTTAAATAATATGCTCTTAAGATTATTTATCAAAATGATATCTTTATTTCAATATTTTACTGGTGTTACCTAATAAATTGGCGACACCATACTGAAACATGATGACTTAAAATATTCAAAATCTAATATATAAGTTGACAGTGCGGCGACACCAAACTTTAAATAAAGTAAATTGTTTTTTAAACTCTTTTTGTTTATTATTTCTTTTTATTTCTCTTTAACACAAAAAATAGAGAGACTTTTTATCAATTAGTCTATAAAGTTTTTTTTACTAAAAAGTATTTTTGTAAAAAAATCGATTTTAACTAGAAATAAATTTTATTTATTTTTAAAAAAATAATTTTTTTGTTCAAATAATTTTTTTTAAAAATTATTCCGACAAGAATAAAAGAATTTGAACAAAAATTATTTTTCAAAAAATAAATAAAATTTATTTTCAATAGGAATCAATTTTTTTACAAAAAGAAAATTTGATAGAAATTTTTTTTGAGTAATTGGTAAGAAGCCTCTCTATTTTTTGTATTAAAAGAAAATAAAAGAAAATAATAAATAGAGAAAGTTTATGAAGCAGTTTAACTTATTTAAAGTTTGATGCTCTCACACTATTAACGGACCTAACAATTCTTTTTAACAACCAATCAAAATGAGAAATTTACGTATTAAATCCCTGAATATTTTAAGTCAATATATTTTAATCTTGTGTCACTAATTTACCCCAACACTGACAAAATACTGTAGTAAAAGTACTATTTTGATAAATAATTTTATGGGTATATATTTGAATTATTAAATTTTTTTATATTATTGATGTAAGAAAAGCCGTATCTTTTATTATTATTACTTCCCACTCTGGCCTCACATGAAAACAAGCAAGGACGGACTCAGTTGATAGTTGAGATGCACAATACACCTCAACTAAAGCGCGTAAGAAACACCTATTCTTCACCGTCGGCCAAGTCTTCGGCATATACCATTAAAATTCCGGTCTTTTATCCACGTGGAAATACAGCGCGTGGATATGGGTCCTACTAAACGATCCAGTCATTCAATCAATATACCCCTTCGAGTACCTTCTCAATGGCAACACAATAAAAATAAAATAAAATAAAATTTCCTTCACAAGGTTAAGCCACGTGTAAAGTGTATTACACGTCAAAAATGACACCCCTCTTCACACGTACGAACGGGCAAAAAAACCACAAAACAGCATAGGCATGGCCTACCTTCTCTTGAATCGAAAAACCAAAGGGGATTCAATGACGAAATTTTAACGTAAGGCATCAACCATTGTGATGATGCCACGTGTATCCTCTCATTATATGTTTCCTCTATAAATAAGCCTCCTAGTCCCAAGCTTTGAA

>Gohir.D03G043600-IPS-Paralogs-Gene-Included

TCCCGCTGGCAGAAAGACCACACGAAGATTTCCAAGTCTGGAGTGCTGATGCGTCGGGTGAGTACACAGTTCGTAGCGCCTATAAACTATTACAAAGTACTGATAATGATCCTAGAGCTTATGCTTTACAAACCAACTATAAGGACTTATACAGGAAACTATGGCCACTTGATCTTCCTTCGAAGATAAAAATTACAGTGTGGAAGATTTCATGGAATTTCCTAGCGACACGTGCTAATATGCTACTCAAAAGGTTAACAAACACATCAGTATGTCCCCGATGTGGCTCAGGAATAGAAACTATAGACCATCTCTTCCGTGAATGTCCTGTTTCAACATCAGGATGGAGAGAGTTATCATTTCATAAGTGTTTGCAAGATAATTAGTTGGGGTTCTTACAGTGGCTAACCTGGGTTTTCTCACAAAACTCGACATCCCAGTGTCGGGTCTTTTGCGTTGCATTATGGGCTATCTGGGGAGACAGGAATAATAAAGTACATAAAAAAGGGAGTAAATCTGGGAAAGAAATAGGGCGATTTGTTCATAGCTACATATCAAAATTAAATGGGATTGGTGAAAACAGACCACTAACTTCCATACCTGTTATAGAATGGAAAAAGCCATCTGACCAAGTTGTTAAGATCAATTTCGACGCAGCATTTGAAGGTAGACAAAATAAAGCGGCTGTTGGAATAGTGGCCATAGACAGAGAGGGTACAGTTATCCTATCATGCTCGGAGGTTTGCCAGAGGGTTCCATCAGCCTTTGCGGCGGAGGCACTCGTATGTTGGAAATCACTCCAGATCGGCGTTTACATGAAGTGGGAAAAAGTTGATATTGAAGGAGATTCTTTATCAATAATAAGGAAATGCAAAGCAAAAAGCCCAGACAAATCAATGGCTAGTGCGTATATTCATGACATCCATCAACTACTGTTAAATACTAAAGATTGCAATTTTGAGCATGTCCCAAGATTAGCAAATAGTCTCGTGCATATCTTAGCAACTGAAGCGTTGAAGAGTAATAAAGGAGATTACCTGATTGGAAGAGTCCCTGAAGTTGCAGAAAGACAAGCGAAAAATGAAAGGTTGAGAGAACCGGATTGAAAGCGGAGAGGGGAGAAATGAAGGGAGGAGAGTTTGGTAAGGGCAGGTAACAGAGTGTCAAAAGCAAAGCCCTTGGATGAAGTGAATTGAAACGTCTTGAAGAAGTAAAGAAAGATGAAAAGACAAGACTTTAATTTTCTTGTAAGGCTGATTGGGTAGGCAGAACAGCAAAATTAATTGCTATAGATTTTGCTTTCTTTTTTTTGGTTTTCTAGACGATTAGGTTTCCTTTTAATTTTGTCTTAGTTTTTTGTTATGTTGGGCCAAGTCTGATTATTAGGCTCTTTATATTTATTTTTAGGACTATATGTGTAGTATTGCAGATTATTTTAATAAAGCCCAATCTGAAATTTTATCAAAAAAATAATAATTGGTTAAAAAATTACTTGATAAGAAAATTTATTATAATTATAAATGAGAATTTAAATTTTCTTAAATGACCTTTTTAAAGGAATTTATATAATAATAAATAAAACTTCAAATTTATATTGAATTTATTTTGCTTCCATTAACCAAACAGTGAAAAGTGAAAAAATTATATTACTTTTTAAATAGTAAATTTTAATTATTTATACAATAATATTTTATATATAAAATTTTATAAATAATACGTTTTTTAATATAATATAATATATTAATTATATATTATCTGTAAAATGTTTTGGGCAATATATTAACGGCTAAGAAAAAATAAAAAATAAAAATTTAAGAATTTAATTTACAATATATCAATTTTTATAATTTAAAAATTACATGAAGTAAAACTACACTATCTATTGAATATGTTAAAAATATTTTAATTATGGTTTCAATTTATTATTATACTATTTAGATTTTAAATAAATTAATATATTTTAATTAAAAATAAAAATTATAAAAATTAAACACATAAAATTACATACAATACATATGACATTAGAAACTAATATATATACTGAAAAATAAATGCTCTTTATTATTTATTATACTTCAAAATTGCCATTAAATTTTTTATTACTAATGAAATTTATCAAAAGTATAACTAAAAATTTTAATTTTATCAAAACGCTTTTTTAATGAATTGAATTTGATAATAGAAAAAAGATGTTTAAAAAAAATTCATTACATACTATATTTATTTTAAGCGCTTTAGAGGATTTTTATTTCTAATAATGTCTAACACATTTAAATTTATCAATAATTTGTTAAGAAATTAATTTAATAAATTTTTTTACATAAAAGTTAAACTTTTTAAAAACCATGTTAATTAAATAATAAGTTTTAAGAGATTATTACTTTTAACATATTTTGAAAAGTTAGCTTCAATATCTAACTAAAACCACCCCACCTTTCATAATTATTGTGAACAATCAAATATATTTTAATATGAAAAAAGGTACATTATAAAAGTTCCAGTGGCATTAAAAAATTGTTAACTACAATACAAAAATAAACTCAAATCTTGATCTCAGTCAAATCCAAATAACTGCCTGCAATTTTAATTTATTTACGACGGGATCTCATCTTTTTAACCAACTCTAAATTTTGGTTTTCTTTATCCTAATAATAATAATAATAATAGCTTAAATAGAATGCCATTCACATCATTTATGAAAATGATATGTTTGCTTCAATGTTTTGTCGGTGCTACTTAACAAATTGGTGACATCAGATTAAAACGTGAGACCTAAAATATTCAGAACTTGATATATAAGCTAAAAGTGTGCCAACACCAAACTTTATATAAAGTAAATTGCTTCAAAAACTCTTCTTGTTTATTATTTCTTTTTATTTCCCTTTAACACAAAATAGAGAGGCCTCTTATTAATTAGTCTATAAAGTTTTCTTCCTACCAAAAAGTATTTTTGTAAAGAAGTCAATTCCGACTAGAAATAAATTTTATATATTTTTTAAAAAATAAATTTTTTGTTCAAATAAATTTTTTAAAATGATTTTTTGAAAAATTATCCCGATAGAAATAAAAGAATTTGAACAAAAAAATTATTTCGAAAAATAAATAAAATTTATTTTCAATTGAAGTCGATTTTTTTTAGAAAAATACATTTTGATAGAAATTTTTTTTGGAGTATTTGGTAAGAGGCCTTTCTATTTTTAGTATTAAAGGAAAATAAAAATAATAATAAATAAAGAGATTTATGAAGCAATTTACCTTATTTAAAGTTTGATACCGCCGCACTATTAACGGATTTAAAACTTTTTTTCAGCAGCCAATCAAAGTGAAAAATTTACATATCAAGTCCTTGTATATTTTAAGTCAACACATTTTAATTTTGTGTCACTGATTTATCAGGTAACACCAATAAAATACTATAGTAAACGTATCATTTTGATAAATAATTTTATGGGTATATCATTTGAATTATTAATTTTTTTTATATTATTGATGTAAGAAAGCCGTAATTTTGTTTTTTTGTTACTTCCCACTCTGGCCTCACGTGAAAACAAGCAAGGACGGACTCAGTTGATAGTTGAGATGCACAATACACCTCAACTAAAGCGCGTAAGAAACATCTATTCTTCACCGTCGGCCAAGCCTTCGGCATATACCATTAAAATACCGGTGTTTTACCCACGTGGACATCAGCGCGTGGATATGGGTCCTACTAGACGATCCAGTCATTCAATCAATATACCCCTTCGAGTACCTTCCCAATGGCAACACAATAAAAATAAAATTTCCTTCACAAGGTTAAGCCACGTGTAAAGTGTATTACACCCCTCTTCACACGTACGAACGGGCAAAAAAACCACAAAACAACAAAGGCATGGCCTACCTTCTCTTCAATCAAAAAACCAAAGGGGATTCAATGACGAAATTTTAACTTAAGGCATCAAACCATTGTGATGATGCCGCGTGTATCCTCTCATTATATGTTTCCTCTATAAAACCTCCTAGTCCCAAGCTTTGAAGCACAAAATTCTGGGGCAAAAAAGAAATAAAAACAGTTTCATTTCCATTCTAAGTTTGTGTTTTTGATTTGCCTTTTGTGGGTTTTTTTTTATAAAAAAGTTCAGCAAAAATGTTTATTGAGAATTTTAAGGTGGAGAGCCCCAATGTGAAGTACACAGAGAATGAAATTCAGTCTGTTTACAACTATGAAACTACAGAGCTTGTTCATGAGAACAAAAATGGAACCTATCAATGGGTTGTTAAACCCAAAACTGTCAAATATGAATTCAAGACTGATATCCATGTCCCTAAATTGGGGTTAGAATCCAAATCTGCCCATCATTTGTTTTTTTGCCATTTCGATTTTGATTATTTTGATTTTATTTATTTATATGTTTTAGGGTGATGCTTGTGGGATGGGGAGGAAACAATGGTTCAACCCTCACCGGTGGTGTTATAGCTAACAAAGAGTGAGTTTTGTACTTTCTAGATAAAAGATCTGATTAATTTTGATTTGTTTTCCTTTCTTTTTTTTTTCTTTTTTTTTTGGTTGATTTTTGAGTTATGGGTAATGATGGGTTTTCTCAGGGGTATCTCTTGGGCTACTAAGGACAAGGTACAACAGGCTAATTACTTTGGTTCATTGACTCAAGCATCAACGATCCGAATTGGGTCTTACAATGGAGAAGAGATTTATGCTCCATTTAAGAGTCTTCTTCCTATGGTATGCGAATTATAAACCCAGATTTAGTTTTTAATTTACTAGTTTGCTACTTAATGCTTAATGCTTGATAATGAATGAATACAGGTGAACCCAAATGATATTGTGTTTGGAGGATGGGACATTAGTGACATGAACCTAGCTGATGCAATGGCTAGGGCCAAGGTTTTCGACATTGATCTGCAAAAGCAACTGAGACCCTACATGGAATCCATGGTCCCACTCCCTGGAATCTACGATCCTGATTTCATTGCTGCTAACCAAGGTGAACGTGCCAACAATGTTATCAAGGGGACCAAGAAAGAACAAGTTCAGCAGGTCATCAAAGACATCAAGTATATAATCAACTCCAACCCTCAAAACTGTATTTGTTAAGCTTGTAGAACAGGAAAAGTAGCTGAATTTTTTAACATTTGATTGTGTAAATCAGGGAGTTCAAAGAGAAAAACAAGGTGGACAAGGTTGTTGTACTCTGGACTGCAAACACTGAGAGGTACAGCAATGTCATCGTGGGGCTAAATGACACCGTGGAAAGCCTTATGGCTTCTTTGGAGAAGAATGAATCAGAGATTTCTCCTTCCACTTTGTATGCTATTGCTTGTGTTCTTGAAAATGTTCCTTTCATCAATGGCAGCCCACAAAACACCTTTGTTCCAGGTTCTTAATTTAGCTCAGCTGTCTTTCTAATTTTATAAATTGTAATGACGAGTAAATATAATAACCTCTACAAGTGACTCTTATTGTATTCAGGGTTGATTGATTTGGCTATTCAAAGGAACTGTCTGATTGGAGGAGATGACTTCAAGAGTGGCCAGACCAAGATGAAATCTGTCCTCGTGGATTTCCTTGTTGGGGCTGGGATCAAGGTATAAATCTTTTAACTTTATGGCACTATCACCATAATTGTTAAAATAAGCAATTAGTACCAATATAATTGAAGTAAAAAGTTTAAGTGTTTGGTAGCTTATGTGTCAATAAAATGTAAATAATATGACATTGATAGGTACTTGTACTAACGGTGATGATAAATTGACAGCCAACATCGATAGTGAGTTACAACCATCTGGGAAATAATGATGGCATGAATCTATCGGCACCCCAAACCTTCCGTTCCAAGGAGATCTCAAAGAGCAATGTTGTTGATGACATGGTTTCAAGCAATGGAATCCTGTATGAGCCTGGTGAACATCCTGATCATGTTGTGGTCATCAAGGTAAATATTTTTTTACGCATCTGTGTCTGTTTGGGCACTTGTCAGTGGTGACCTTAAAGCTTTTTTGCTTGATTGGTCTTAAAATGTGTACTTTGTCTTCCTTGTATTATAGTATGTGCCATATGTGGGAGACAGCAAGAGAGCCATGGATGAGTATACATCAGAGATATTCATGGGAGGCAAGAACACCATTGTGTTGCACAACACATGTGAGGATTCCCTGTTGGCTGCTCCCATTATCCTAGATTTGGTTCTCCTTGCTGAGCTTAGCACCAGGATCCAGTTCAAGGCTGATGGAGAGGTATGACCCTTAGAACTTTGTGGACTCTTCTGGGTTGACCCTTAAAACATATTTTTGCAAAAACATTGATGCTGAATTTTGGTTTTGCAGGGCAAGTTCCACTCTTTCCATCCTGTGGCTACAATCCTCAGTTACCTCACCAAAGCCCCTCTTGTGAGTCCAAAGCATCCCTTCGTCTTTTCCTGTAACTCTGGCCTACTAAAAATATAATAATAATTAATAATAAAACAGGGGTGGGAAGGGTGACACCTTCACCAATAAAACATTGATATTATTGTCTTGAATTAACTGGTAACTCAACTGTTTATAGTTTAAGTGGGAAACCAGGAATCGCAAATAATTAATAATAAAATGGGGGTGGAGAGGGGTGACACCTTCACCAATAAAACATTGATATTATTGTCTTGAATTAAATGGTAACTCAACTGTTTATAGTTTAAGTGGGAAACAAGGAATCGCAAATTCACTTTATGGTGCCTCTTTGTGATATATTTCATGTGACCATCAATGATTTTATGTAAAGCTGCTTCTGTCTATTTCACATGAAGCAACTGGTGAGAGGGGACATAATATTGAATTAAAGTTTGTGAACTTTATGTCAAATTTCAGTTCAAGAATGTGAATAGAATCATGACAATCACTTGAACCAGAAATCTGTGAAGAAATTTACTGAGAGCTGACATTGTTTTGGCAAAAAACAGGTTCCACCAGGCACACCGGTGGTGAACGCACTGTCCAAGCAGCGTGCAATGCTGGAGAACATACTAAGGGCCAGCATTGGCTTGGCTCCTGAAAACAACATGATTTTGGAATACAAGTGAAGAAGTTGAGGAGAGAAAAGTGGAACAAAGCTTTGGTTCTTAGAACTTTAGGACCATTTTCCCTTTCTCTGTCGTATTCATAAAAGCTGTGGTTACTGTAGTCACAGATTGTAACGGTTGTATGATGGTTTATGCTTTGGTATATTATGCTTCTTTTTTTCTCATTTATTTTCCGATGGCGCTCATGTTTGTAGCTCACTCTTGAACCAAGCACCCCTTTTGTGGAAATAGATATGATAATATGAATTTTGGTAACCGTGTTATC

>Gohir.D03G043600-IPS-Paralogs-promoterOnly

TCCCGCTGGCAGAAAGACCACACGAAGATTTCCAAGTCTGGAGTGCTGATGCGTCGGGTGAGTACACAGTTCGTAGCGCCTATAAACTATTACAAAGTACTGATAATGATCCTAGAGCTTATGCTTTACAAACCAACTATAAGGACTTATACAGGAAACTATGGCCACTTGATCTTCCTTCGAAGATAAAAATTACAGTGTGGAAGATTTCATGGAATTTCCTAGCGACACGTGCTAATATGCTACTCAAAAGGTTAACAAACACATCAGTATGTCCCCGATGTGGCTCAGGAATAGAAACTATAGACCATCTCTTCCGTGAATGTCCTGTTTCAACATCAGGATGGAGAGAGTTATCATTTCATAAGTGTTTGCAAGATAATTAGTTGGGGTTCTTACAGTGGCTAACCTGGGTTTTCTCACAAAACTCGACATCCCAGTGTCGGGTCTTTTGCGTTGCATTATGGGCTATCTGGGGAGACAGGAATAATAAAGTACATAAAAAAGGGAGTAAATCTGGGAAAGAAATAGGGCGATTTGTTCATAGCTACATATCAAAATTAAATGGGATTGGTGAAAACAGACCACTAACTTCCATACCTGTTATAGAATGGAAAAAGCCATCTGACCAAGTTGTTAAGATCAATTTCGACGCAGCATTTGAAGGTAGACAAAATAAAGCGGCTGTTGGAATAGTGGCCATAGACAGAGAGGGTACAGTTATCCTATCATGCTCGGAGGTTTGCCAGAGGGTTCCATCAGCCTTTGCGGCGGAGGCACTCGTATGTTGGAAATCACTCCAGATCGGCGTTTACATGAAGTGGGAAAAAGTTGATATTGAAGGAGATTCTTTATCAATAATAAGGAAATGCAAAGCAAAAAGCCCAGACAAATCAATGGCTAGTGCGTATATTCATGACATCCATCAACTACTGTTAAATACTAAAGATTGCAATTTTGAGCATGTCCCAAGATTAGCAAATAGTCTCGTGCATATCTTAGCAACTGAAGCGTTGAAGAGTAATAAAGGAGATTACCTGATTGGAAGAGTCCCTGAAGTTGCAGAAAGACAAGCGAAAAATGAAAGGTTGAGAGAACCGGATTGAAAGCGGAGAGGGGAGAAATGAAGGGAGGAGAGTTTGGTAAGGGCAGGTAACAGAGTGTCAAAAGCAAAGCCCTTGGATGAAGTGAATTGAAACGTCTTGAAGAAGTAAAGAAAGATGAAAAGACAAGACTTTAATTTTCTTGTAAGGCTGATTGGGTAGGCAGAACAGCAAAATTAATTGCTATAGATTTTGCTTTCTTTTTTTTGGTTTTCTAGACGATTAGGTTTCCTTTTAATTTTGTCTTAGTTTTTTGTTATGTTGGGCCAAGTCTGATTATTAGGCTCTTTATATTTATTTTTAGGACTATATGTGTAGTATTGCAGATTATTTTAATAAAGCCCAATCTGAAATTTTATCAAAAAAATAATAATTGGTTAAAAAATTACTTGATAAGAAAATTTATTATAATTATAAATGAGAATTTAAATTTTCTTAAATGACCTTTTTAAAGGAATTTATATAATAATAAATAAAACTTCAAATTTATATTGAATTTATTTTGCTTCCATTAACCAAACAGTGAAAAGTGAAAAAATTATATTACTTTTTAAATAGTAAATTTTAATTATTTATACAATAATATTTTATATATAAAATTTTATAAATAATACGTTTTTTAATATAATATAATATATTAATTATATATTATCTGTAAAATGTTTTGGGCAATATATTAACGGCTAAGAAAAAATAAAAAATAAAAATTTAAGAATTTAATTTACAATATATCAATTTTTATAATTTAAAAATTACATGAAGTAAAACTACACTATCTATTGAATATGTTAAAAATATTTTAATTATGGTTTCAATTTATTATTATACTATTTAGATTTTAAATAAATTAATATATTTTAATTAAAAATAAAAATTATAAAAATTAAACACATAAAATTACATACAATACATATGACATTAGAAACTAATATATATACTGAAAAATAAATGCTCTTTATTATTTATTATACTTCAAAATTGCCATTAAATTTTTTATTACTAATGAAATTTATCAAAAGTATAACTAAAAATTTTAATTTTATCAAAACGCTTTTTTAATGAATTGAATTTGATAATAGAAAAAAGATGTTTAAAAAAAATTCATTACATACTATATTTATTTTAAGCGCTTTAGAGGATTTTTATTTCTAATAATGTCTAACACATTTAAATTTATCAATAATTTGTTAAGAAATTAATTTAATAAATTTTTTTACATAAAAGTTAAACTTTTTAAAAACCATGTTAATTAAATAATAAGTTTTAAGAGATTATTACTTTTAACATATTTTGAAAAGTTAGCTTCAATATCTAACTAAAACCACCCCACCTTTCATAATTATTGTGAACAATCAAATATATTTTAATATGAAAAAAGGTACATTATAAAAGTTCCAGTGGCATTAAAAAATTGTTAACTACAATACAAAAATAAACTCAAATCTTGATCTCAGTCAAATCCAAATAACTGCCTGCAATTTTAATTTATTTACGACGGGATCTCATCTTTTTAACCAACTCTAAATTTTGGTTTTCTTTATCCTAATAATAATAATAATAATAGCTTAAATAGAATGCCATTCACATCATTTATGAAAATGATATGTTTGCTTCAATGTTTTGTCGGTGCTACTTAACAAATTGGTGACATCAGATTAAAACGTGAGACCTAAAATATTCAGAACTTGATATATAAGCTAAAAGTGTGCCAACACCAAACTTTATATAAAGTAAATTGCTTCAAAAACTCTTCTTGTTTATTATTTCTTTTTATTTCCCTTTAACACAAAATAGAGAGGCCTCTTATTAATTAGTCTATAAAGTTTTCTTCCTACCAAAAAGTATTTTTGTAAAGAAGTCAATTCCGACTAGAAATAAATTTTATATATTTTTTAAAAAATAAATTTTTTGTTCAAATAAATTTTTTAAAATGATTTTTTGAAAAATTATCCCGATAGAAATAAAAGAATTTGAACAAAAAAATTATTTCGAAAAATAAATAAAATTTATTTTCAATTGAAGTCGATTTTTTTTAGAAAAATACATTTTGATAGAAATTTTTTTTGGAGTATTTGGTAAGAGGCCTTTCTATTTTTAGTATTAAAGGAAAATAAAAATAATAATAAATAAAGAGATTTATGAAGCAATTTACCTTATTTAAAGTTTGATACCGCCGCACTATTAACGGATTTAAAACTTTTTTTCAGCAGCCAATCAAAGTGAAAAATTTACATATCAAGTCCTTGTATATTTTAAGTCAACACATTTTAATTTTGTGTCACTGATTTATCAGGTAACACCAATAAAATACTATAGTAAACGTATCATTTTGATAAATAATTTTATGGGTATATCATTTGAATTATTAATTTTTTTTATATTATTGATGTAAGAAAGCCGTAATTTTGTTTTTTTGTTACTTCCCACTCTGGCCTCACGTGAAAACAAGCAAGGACGGACTCAGTTGATAGTTGAGATGCACAATACACCTCAACTAAAGCGCGTAAGAAACATCTATTCTTCACCGTCGGCCAAGCCTTCGGCATATACCATTAAAATACCGGTGTTTTACCCACGTGGACATCAGCGCGTGGATATGGGTCCTACTAGACGATCCAGTCATTCAATCAATATACCCCTTCGAGTACCTTCCCAATGGCAACACAATAAAAATAAAATTTCCTTCACAAGGTTAAGCCACGTGTAAAGTGTATTACACCCCTCTTCACACGTACGAACGGGCAAAAAAACCACAAAACAACAAAGGCATGGCCTACCTTCTCTTCAATCAAAAAACCAAAGGGGATTCAATGACGAAATTTTAACTTAAGGCATCAAACCATTGTGATGATGCCGCGTGTATCCTCTCATTATATGTTTCCTCTATAAAACCTCC
